# Associations between floor material and *E. coli* contamination in rural Bangladeshi households

**DOI:** 10.1101/2025.05.07.652756

**Authors:** Sumaiya Tazin, Mahfuza Islam, Amy J Pickering, Laura H Kwong, Andrew Mertens, Caitlin Niven, Benjamin F Arnold, Alan E Hubbard, Mahfuja Alam, Debashis Sen, Sharmin Islam, Mahbubur Rahman, Leanne Unicomb, Stephen P Luby, John M Colford, Jade Benjamin-Chung, Ayse Ercumen

## Abstract

Soil floors are common in low-income countries and can harbor contamination from unsafely managed human and animal fecal waste. Soil/dust ingestion directly from floors or indirectly via hands, drinking water and food can significantly contribute to children’s ingestion of fecal organisms. We assessed if finished (e.g., concrete) floors are associated with lower *E. coli* contamination in the domestic environment in rural Bangladesh. We collected samples from 1864 households over 3.5 years, including stored drinking water, child and caregiver hand rinses, courtyard soil, food, and flies (n=24,118 samples), and enumerated *E. coli* using IDEXX Quanti-Tray/2000. Controlling for potential confounders (socio-demographics, water/sanitation status, animal ownership), households with finished floors had slightly lower log10-transformed *E. coli* counts (Δlog10= -0.10 (-0.20, 0.00)) and prevalence (prevalence ratio=0.90 (0.83, 0.98)) on child hands than households with soil floors; floor material was not associated with contamination levels in other sample types. Finished floors were associated with lower *E. coli* contamination of child hands, food and stored drinking water following periods of higher rainfall and temperature, and lower *E. coli* contamination of child hands in households with more domestic animals. Measures to control enteric infections in low-income countries should test flooring improvements to reduce exposure to fecal contamination.

## Introduction

In settings where fecal waste is not well contained, children are frequently exposed to pathogens and experience a high burden of enteric infections and growth faltering. Enteropathogens are transmitted from the feces of infected humans or animals to the gastrointestinal tract of new hosts through various environmental pathways. In particular, fecally contaminated soil in the domestic environment is increasingly recognized as a major source of pathogen exposure for young children in low-resource settings. Children have frequent hand and object contact with soil and can accidentally or deliberately ingest soil and/or dust,^1, 2^ contributing substantially to their ingestion of fecal organisms.^3^ Soil ingestion has been associated with increased risk of diarrhea, environmental enteric dysfunction and stunting in children.^4–8^

Many homes in low-income countries have floors made of soil,^9^ and children may play, eat and sleep on soil floors, presenting a potentially substantial soilborne exposure pathway. Soil floors in low-income countries can harbor soil-transmitted helminths and other parasites, including hookworm,^10^ *Ascaris lumbricoides,*^11–13^ *Strongyloides stercoralis,*^14^ and *Giardia duodenalis*.^12–15^ The eggs of some soil-transmitted helminths (e.g., *A. lumbricoides*) can remain infectious in soil for several months under favorable conditions, perpetuating infections.^16^ Studies have also detected high levels of fecal indicator bacteria on interior and exterior soil floors,^17–20^ and fecal contamination of domestic soil has been associated with increased contamination along other fecal-oral transmission pathways, including child hands, stored water and food.^21^

A finished (e.g., concrete) floor can reduce hand and object contact with indoor soil and accidental and/or deliberate soil ingestion by young children, as well as provide a durable cleanable surface that does not support pathogen survival. Interventions to improve housing conditions have been associated with improved health outcomes, including respiratory and mental health.^22^ Specifically, having finished floor materials as opposed to soil floors has been associated with a range of health benefits in children. A 2023 meta-analysis concluded that soil household floors are a potential contributor to enteric and parasitic infections in low- and middle-income countries.^25^ Replacing soil floors with concrete floors in a large-scale Mexican program reduced diarrhea by 13% and parasite infections in children under six years by 20%.^23^ In Zimbabwe, improved household flooring reduced infant diarrheal illness irrespective of the quality of the household’s water or sanitation.^24^ However, none of these studies investigated the environmental pathways behind these health improvements to substantiate the observed health effects associated with improved floors and explore what mechanism drives them.

Here, we use data from a randomized controlled trial in Bangladesh (WASH Benefits) to assess associations between floor material and fecal indicator bacteria measured along several pathways in the household environment. The WASH Benefits trials assessed the effects of individual and combined water quality, sanitation, handwashing (WASH) and nutrition interventions on child health in rural Kenya and Bangladesh. In Bangladesh, the WASH interventions reduced child diarrhea and enteric infections by 31-40%,^26–29^ but did not improve child growth.^26^ In Kenya, the WASH intervention reduced *A. lumbricoides* infections by 22%,^34^ but had no effect on *Giardia* infections, diarrhea, and child growth.^34,35^ In both countries, the WASH interventions achieved limited reductions in environmental contamination with fecal indicators and pathogens, and the domestic environment remained heavily contaminated despite the interventions.^30–33, 36^ These findings prompted calls for “transformative” interventions that more effectively interrupt infectious disease transmission.^37–38^

An observational analysis of child health data from both the WASH Benefits Bangladesh and Kenya trials showed that children in homes with finished floors had 38-67% lower prevalence of soil-transmitted helminths and 18-22% lower prevalence of *Giardia duodenalis* infections compared to children in homes with soil floors.^39^ To examine the environmental mechanisms behind these associations, here we use data from a subset of households enrolled in the WASH Benefits Bangladesh trial to assess associations between finished vs. soil floors and fecal contamination along various fecal-oral transmission pathways, including child and caregiver hands, stored drinking water, food, courtyard soil and flies.

## Methods

### Trial design

WASH Benefits included two cluster-randomized trials that provided WASH and child nutrition interventions -alone and in combination- to rural households in Kenya and Bangladesh.^40^ In Bangladesh, participants were selected from the rural villages of Gazipur, Kishoreganj, Mymensingh, and Tangail districts in central Bangladesh. Households where pregnant women in their first or second trimester lived (N=5551) were selected for enrollment. Groups of 6-8 proximate enrolled households were organized into clusters, and eight adjacent clusters were further grouped into geographic blocks. Within each block, clusters were randomized into intervention vs. control groups. The trial design and specifics of the interventions have been previously reported.^40^

### Assessment of floor material

Household characteristics, including floor material, were recorded at enrollment and at one and two years after intervention initiation when the trial visited households to record child health outcomes. Field staff observed the main floor material of the room where the enrolled pregnant woman or the index child (child who was in utero at enrollment) slept. Floors were considered finished if the most common floor material was wood, tile, or concrete, and unfinished if the floor material was entirely or mostly made of soil.

### Environmental sub-studies

Environmental contamination measures were pre-specified as intermediate outcomes of the WASH Benefits trial. We use data from two sub-studies that measured environmental contamination among a subset of households enrolled in the trial.^31, 33^ The first sub-study was conducted on average 4 months after the initiation of interventions and enrolled a total of 1840 households from the control, sanitation, and combined WASH arms of the trial for a cross-sectional assessment. ^31^ Samples were collected from stored drinking water, food given to young children, hand rinses from index children, courtyard soil, flies captured near the kitchen, groundwater from tubewells, and ponds adjacent to enrolled households. The second sub-study was conducted between 1 and 3.5 years after the initiation of interventions and enrolled a total of 720 households from the control and sanitation arms of the trial for eight longitudinal assessments. Samples were collected approximately quarterly from stored drinking water, food given to young children, hand rinses from index children and their caregivers, and courtyard soil. We combined data from these two sub-studies, resulting in nine rounds of environmental measurements from 1864 unique households over approximately 3.5 years. The present analysis is focused on samples of stored drinking water, food, index child and caregiver hand rinses, courtyard soil and flies. We excluded tubewell and pond samples from our analysis because we did not expect indoor floor materials to affect contamination in these sample types.

#### Sample collection

To collect stored drinking water, field staff asked participants to supply a glass of water from their storage container the same way they would give it to a child <5 years old. Participants were instructed to pour the water into a sterile Whirlpak bag (Nasco, Modesto, CA) to collect approximately 150 mL of sample. If respondents reported using chlorine to treat their water, sodium thiosulfate was utilized to neutralize any residual chlorine. Food samples were collected by asking participants to provide a small portion of solid stored food in the same manner they feed their children, with a preference for rice. The food was scooped to fill a 50 mL sterile plastic tube using a sterile spoon. Hand rinses were collected from both index children and their caregivers. A hand rinse was collected from the youngest child <5 years if the index child was unavailable. Sampled individuals placed one hand at a time into a sterile Whirlpak bag pre-filled with 250 mL of distilled water. The hand was massaged from outside the bag for 15 seconds and shaken for 15 seconds. The same procedure was repeated for the other hand within the same bag, and the rinse water was preserved. Soil samples were obtained from a 30×30 cm area in the courtyard at the entrance of the enrolled household. Field workers designated the area using an alcohol-sterilized metal stencil and used a sterile plastic scoop to scrape the top layer of soil within the stencil once vertically and once horizontally into a sterile Whirlpak bag to obtain approximately 50 g of soil. To collect flies, field workers horizontally hung three 1.5-foot strips of non-baited sticky fly tape at suitable location (away from stove smoke, protected from rain, typically under a roof) in the kitchen and latrine areas. After 3-6 hours, field workers returned to the household and removed one fly from the center of the strip with the highest number of flies using alcohol-sterilized tweezers and placed it into a sterile Whirlpak bag. Field workers collected 10% field blanks for stored drinking water and hand rinse samples. For water samples, blanks were collected by transferring 100 mL of sterile water into the sterile Whirlpak in the same manner the water samples were collected. For hand rinse samples, blanks were collected by opening the pre-filled Whirlpak and performing the massaging and shaking steps without immersing a child or caregiver hand in the Whirlpak. No blanks were collected for food, soil, and fly samples. All samples were transported on ice to the field laboratory of the International Centre for Diarrhoeal Disease Research, Bangladesh (icddr,b) and processed within 12 hours of collection.

#### Sample processing

Samples were analyzed using IDEXX Quanti-Tray/2000 to enumerate the most probable number (MPN) of *E. coli*. 100 mL aliquots of stored water samples were analyzed without dilution. For hand rinse samples, 50 mL of sample was diluted with 50 mL of distilled water. Food and soil samples were homogenized with distilled water in a sterile blending bag using a laboratory-scale food processor. For food samples, a 10 g aliquot was homogenized with 100 mL of water. From the resulting homogenate, 10 mL was mixed with 90 mL of distilled water to produce a 100 mL aliquot. For soil samples, a 20 g aliquot was homogenized with 200 mL of water. Then, 1 mL of the resulting homogenate was mixed with 99 mL of distilled water, and 1 mL of this solution was further diluted with 99 mL of distilled water. The moisture content of food and soil samples was determined by oven-drying 5 g aliquots at 110 °C for 24 hours. Flies were homogenized with a pestle from outside the Whirlpak and mixed with 100 mL of distilled water to make a slurry. 1 mL of this slurry was then mixed with 99 mL of distilled water. Laboratory staff processed 5% of samples in replicate and performed 10% laboratory blanks. Colilert-18 media was added to the 100-mL sample aliquots. Trays were incubated at 44.5 °C for 18 hours.^41^ Bacterial counts obtained were expressed in MPN per 100 mL for water samples, per 2 hands for child and caregiver hand rinses, per 1 dry gram for food and soil samples, and per 1 fly for flies.

### Ethics

Participants provided written informed consent in the local language (Bengali). The study protocol was reviewed and approved by human subjects committees at the icddr,b (PR-11063), University of California, Berkeley (2011-09-3652), and Stanford University (25863).

### Statistical methods

We conducted a prospective, observational analysis of associations between household floor material and subsequently measured environmental contamination. We used the floor material recorded at baseline for our primary analysis as this assessment preceded all environmental measurements. To account for changes in floor material over time, we conducted a sensitivity analysis where we only used data from households that had the same floor material at both enrollment and the two-year follow-up. Our outcomes of interest were the prevalence and abundance of *E. coli* in the environmental samples. We quantified abundance using log10-transformed MPN counts after replacing non-detects with half the lower detection limit (0.5 MPN) and values above the upper detection limit of 2419.6 MPN with 2420 MPN.

#### Estimation strategy

We estimated log10-transformed *E. coli* count differences (Δlog10) and prevalence ratios (PR) to compare households with finished vs. unfinished floors, separately for each sample type and pooling data across all trial arms and data collection rounds. We used generalized linear models with robust standard errors at the study block level to account for cluster-randomization and repeated measurements. Models controlled for potential confounders, which were measured either at the trial’s baseline (caregiver’s age and education, number of children <18 years in the household, number of individuals living in the compound, food security (assessed by the Household Food Insecurity Access Scale (HFIAS) index),^42^ asset-based wealth index, wall materials, drinking water source, minutes to primary drinking water source, and number of cows, goats, and chickens) or at the time of sample collection (sex and age of index child). Models for food and stored water samples included additional variables measured at the time of sample collection. For water samples, we controlled for whether the storage container was covered and had a narrow mouth and the number of hours the water had been stored. For food samples, we controlled for the number of hours since the food was prepared. All models also controlled for study arm (binary variable for intervention vs. control).

#### Effect modification

We assessed potential modification of the association between floor material and *E. coli* contamination by rainfall, temperature, and animal ownership. We used publicly available daily observations from GloH2O’s Multi-Source Weighted-Ensemble Precipitation (MSWEP) dataset and the National Aeronautics and Space Administration’s Famine and Early Warning Systems Network (FEWS NET) Land Data Assimilation System for Central Asia (FLDAS-Central Asia) dataset and matched daily rainfall and temperature values to our study data by date and GPS coordinate.^33^ We used four binary effect modification variables: two indicators for whether heavy rain and elevated temperature occurred within 2 days prior to sample collection, and two indicators for whether the 2-day rolling-average rainfall or temperature prior to sample collection were above median. Heavy rainfall and elevated temperature were defined as >80^th^ percentile of daily values for the study region over the 3.5-year data collection period.^43, 44^ We also generated a categorical effect modification variable based on tertiles of the total number of animals owned by the households. We estimated associations between floor material and *E. coli* outcomes within each rainfall, temperature and animal ownership subgroup and included interaction terms between floor material and the effect modification variables. A p-value <0.20 on the interaction term was interpreted as significant effect modification. The effect modification models controlled for the same covariates as the primary models.

## Results

### Enrollment

The first environmental sub-study visited households once between July 2013 to March 2014, and the second sub-study visited households eight times between June 2014 and December 2016 to collect a total of 6350 stored drinking water samples, 2181 stored food samples, 7092 child hand rinse samples, 5397 caregiver hand rinse samples, 2538 courtyard soil samples, and 610 fly samples (Table S1).

### Floor material

Among 1864 unique households with available environmental samples, 11.8% (219) had finished floors and 88.3% (1645) had unfinished floors, as measured at the baseline of the trial. Of 1864 households, 91.3% (1703) had available floor material data from the two-year follow-up. Among these 1703 households, 93.8% (1597) retained the same floor material at the two-year follow-up, while 4.3% (74) transitioned from soil to concrete floors and 1.9% (32) from concrete to soil floors. Of the 1597 households that retained the same floor material, 10.8% (173) had finished floors and 89.2% (1424) had unfinished floors at both measurement timepoints.

Households with finished floors generally had higher levels of education and were wealthier and more food secure. On average, 83.1% of caregivers in households with finished floors at baseline had secondary or higher education compared to 49.8% in households with unfinished floors (Table 1). Among households with finished floors, 93.6% reported being food secure compared to 64.9% of households with unfinished floors (Table 1). Households with finished floors were more likely to have electricity, improved wall materials and assets (e.g., television, mobile phone, bicycle, motorcycle) while households with unfinished floors appeared to own slightly more animals (Table 1).

**Table 1.**
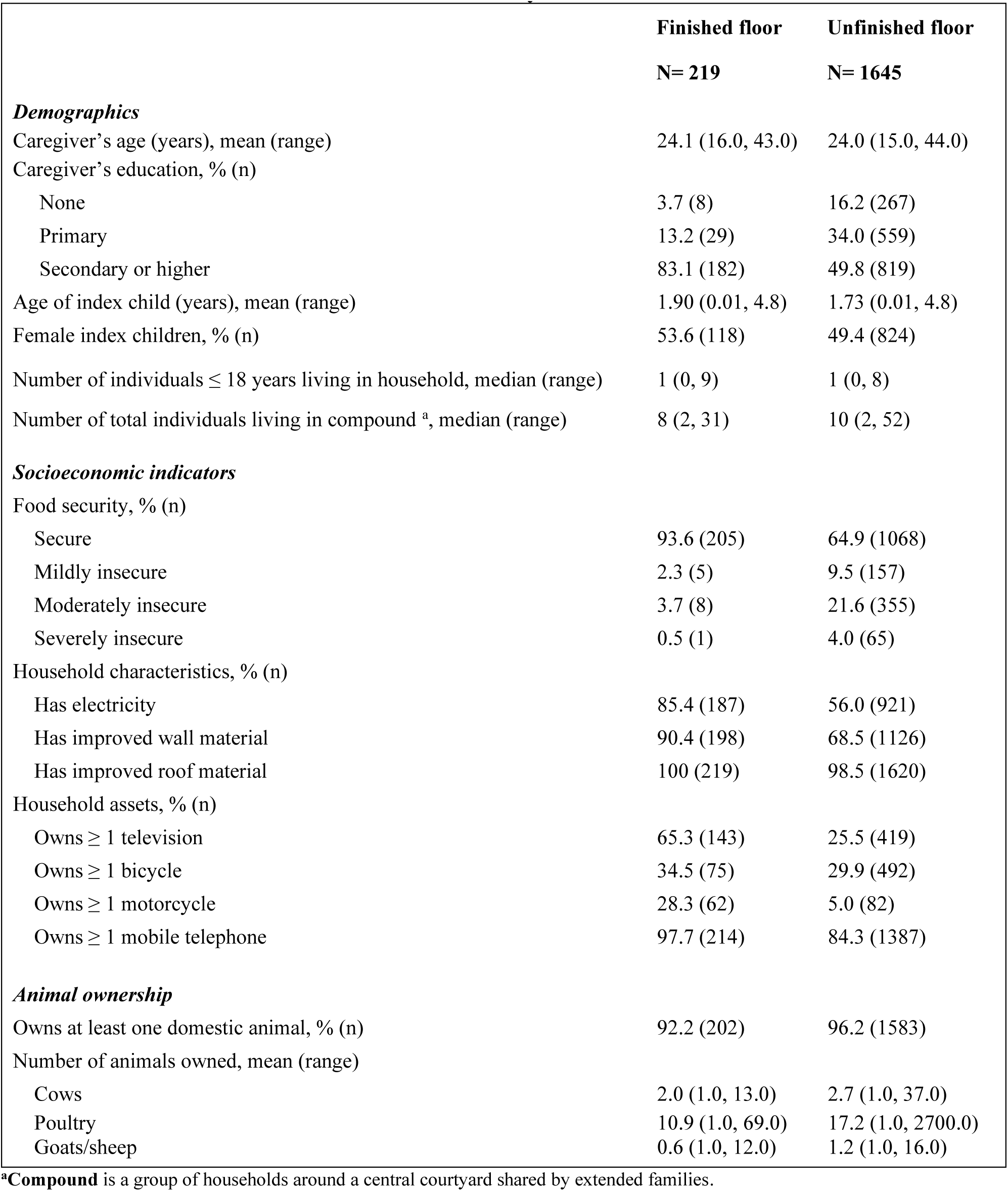
Characteristics of enrolled households by floor material.

### E. coli counts and prevalence

In households with unfinished floors, log-10 transformed counts of *E. coli* ranged from 0.7 in food samples to 5.1 in courtyard soil samples (Fig 1, Table S2), while in households with finished floors, log-10 transformed counts ranged from 0.6 in food samples to 5.0 in courtyard soil samples (Fig 1, Table S2). In households with unfinished floors, *E. coli* prevalence ranged from 54% in fly samples to 95% in courtyard soil samples (Fig S1, Table S3), and in households with finished floors, *E. coli* prevalence ranged from 52% in fly samples to 93% in courtyard soil samples (Fig S1, Table S3). For all sample types, households with finished floors had slightly lower *E. coli* counts and prevalence than those with unfinished floors (Fig 1, Fig S1).

**Figure 1:**
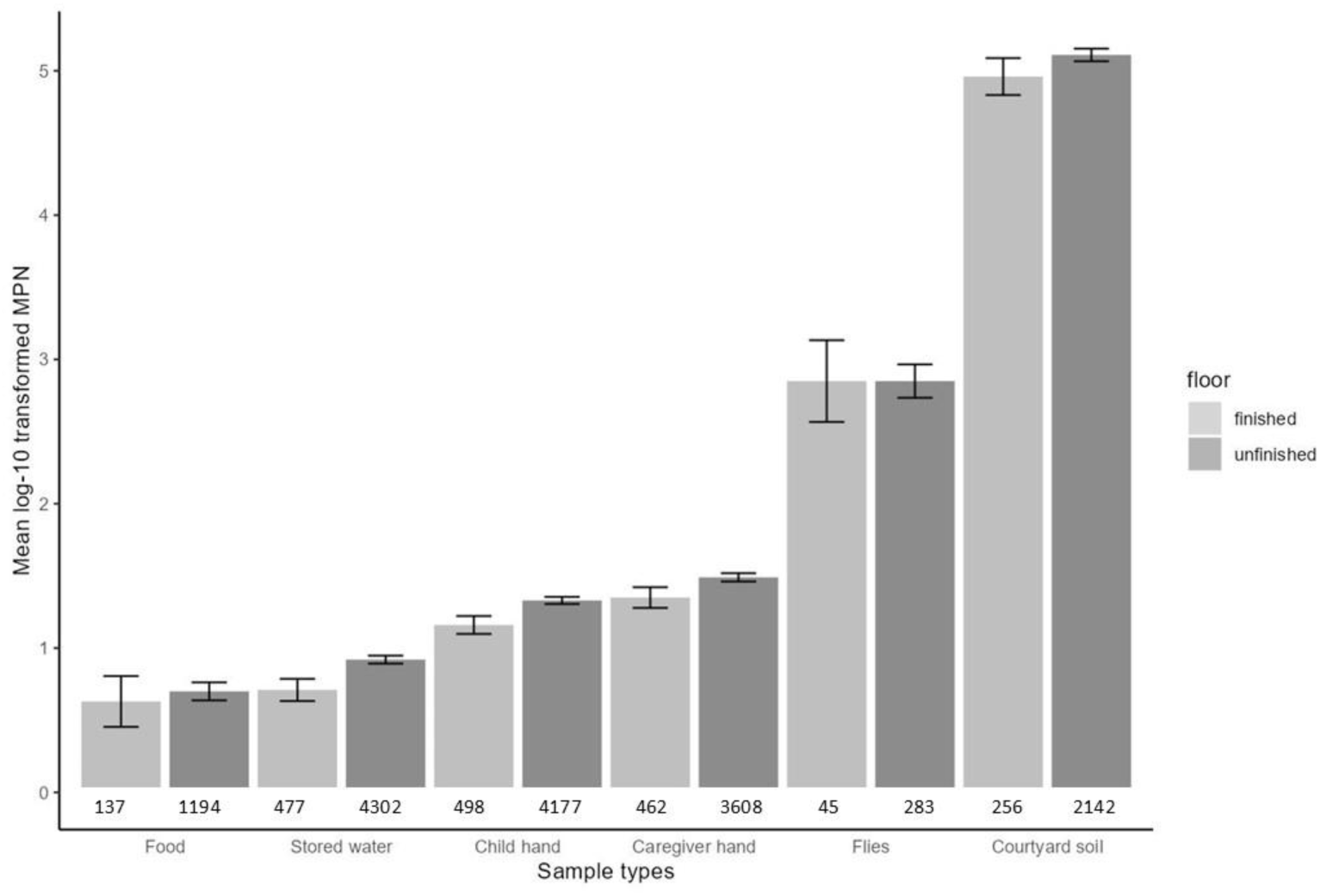
Mean log10-transformed most probable number (MPN) of *E. coli* with 95% confidence intervals (CI) by sample type and floor material. Each pair of columns represents a sample type. The number of samples is listed below the columns. Units by sample type are per 100 mL of stored water, two child and caregiver hands, one dry gram of soil and food, and one fly.

### Associations between floor material and E. coli counts and prevalence

In unadjusted analyses, having a finished floor was associated with associated with lower mean log10-transformed *E. coli* counts in stored drinking water (Δlog10 = −0.21, (−0.31, −0.11), child hand rinses (Δlog10 = −0.17 (−0.26, −0.09)), and caregiver hand rinses (Δlog10 = −0.13 (−0.23, −0.04)) (Fig 2, Table S2). In adjusted analyses, having a finished floor was associated with slightly lower log10-transformed *E. coli* counts (Δlog10 = −0.10 (−0.20, 0.00)) on child hands only (Fig 2, Tables S2). Findings were similar for *E. col*i prevalence. In unadjusted analyses, having a finished floor was associated with 6-13% lower *E. coli* prevalence in stored drinking water, and caregiver and child hand rinses, while in adjusted analyses, only child hand rinse samples had slightly (10%) lower *E. coli* prevalence (Fig S2, Table S3). In the sensitivity analysis limited to households who had the same floor material at baseline and the two-year follow-up, point estimates for associations between floor material and *E. coli* counts and prevalence on child hands remained similar to the primary analysis but could no longer be distinguished from chance, and there was no statistically significant association between floor material and *E. coli* counts or prevalence for any other sample type (Tables S4 and S5).

**Figure 2:**
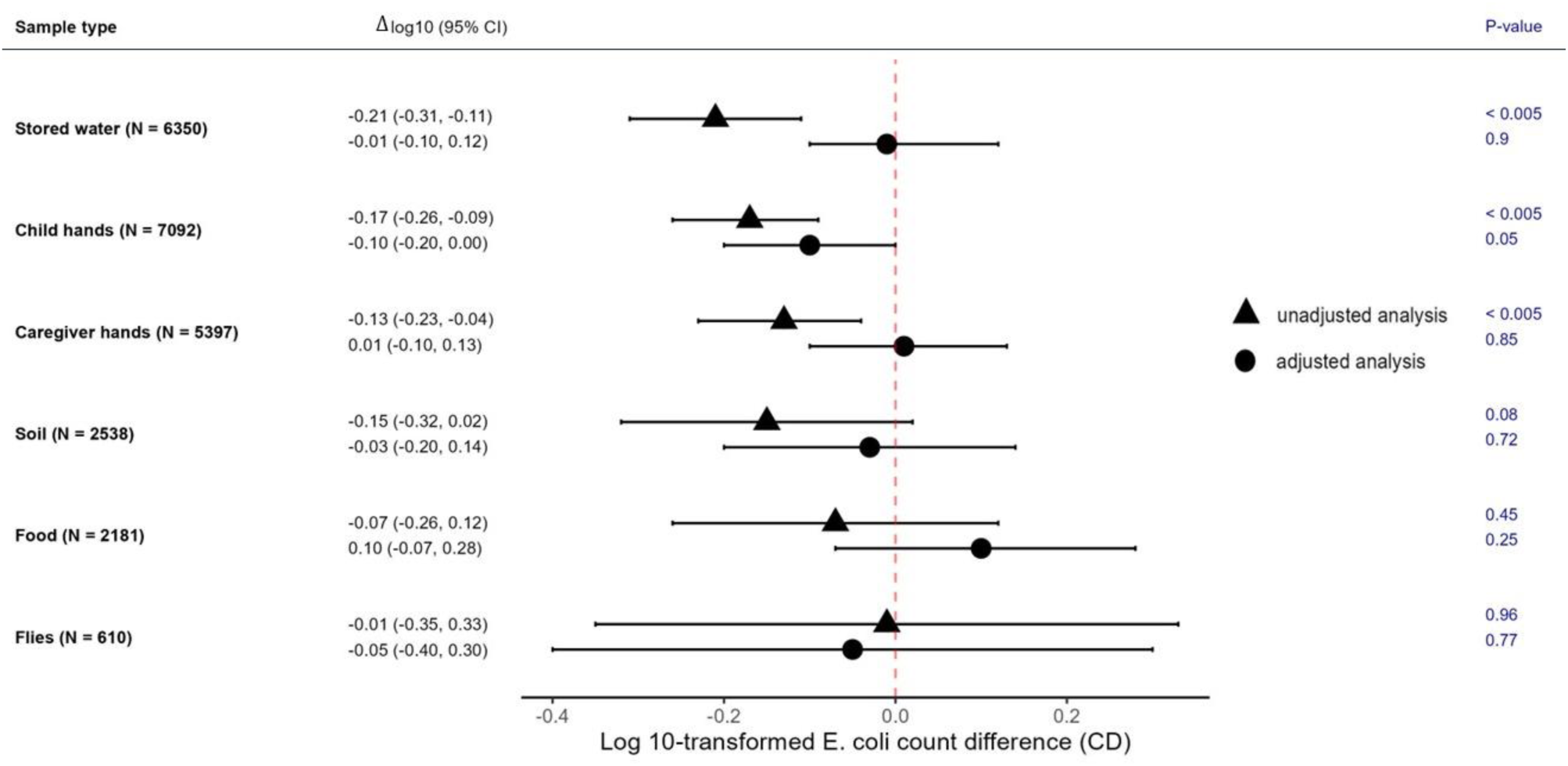
Associations between floor material and log10-transformed *E. coli* counts in environmental samples by sample type. Unadjusted and adjusted log10-transformed count differences (Δlog10) for finished vs unfinished floors, 95% confidence intervals (CIs) and associated p-values are shown for each sample. Adjusted analyses controlled for study arm (intervention vs. control), sex and age of index child, caregiver’s age and education, number of children <18 years in the household, number of individuals living in the compound, food security, asset-based wealth index, wall materials, drinking water source, minutes to the primary drinking water source, number of cows, chickens and sheep/goats, narrow mouth container (only for stored water samples), cover status of the container (only for stored water samples), hours since water has been stored (only for stored water samples), and hours since stored food was prepared (only for food samples). Units by sample type are per 100 mL of stored water, two child and caregiver hands, one dry gram of soil and food, and one fly.

### Effect modification by rainfall and temperature

Following heavy rain, above-median rain and above-median temperature, finished floors were associated with 0.14 to 0.23 log lower *E. coli* counts on child hands (interaction p-values<0.20 compared to periods of no heavy rain, below-median rain and below-median temperature, Fig 3, Table S6). Following heavy/above-median rain and elevated/above-median temperature, finished floors also appeared to be associated with 0.11 to 0.32 log lower *E. coli* counts in food (interaction p-values<0.20 compared to periods of no heavy rain, below-median rain, no elevated temperature and below-median temperature, Fig 3, Table S6). However, the associations between floor material and *E. coli* counts in food could not be distinguished from chance in any rainfall or temperature stratum (Fig 3, Table S6). Following periods of above-median temperature, finished floors were associated with 0.60 log lower *E. coli* counts in/on flies (Fig 3, Tables S6). There was limited evidence of effect modification by rainfall or temperature on *E. coli* counts in stored water, on caregiver hands and in courtyard soil (Fig 3, Table S6).

**Figure 3:**
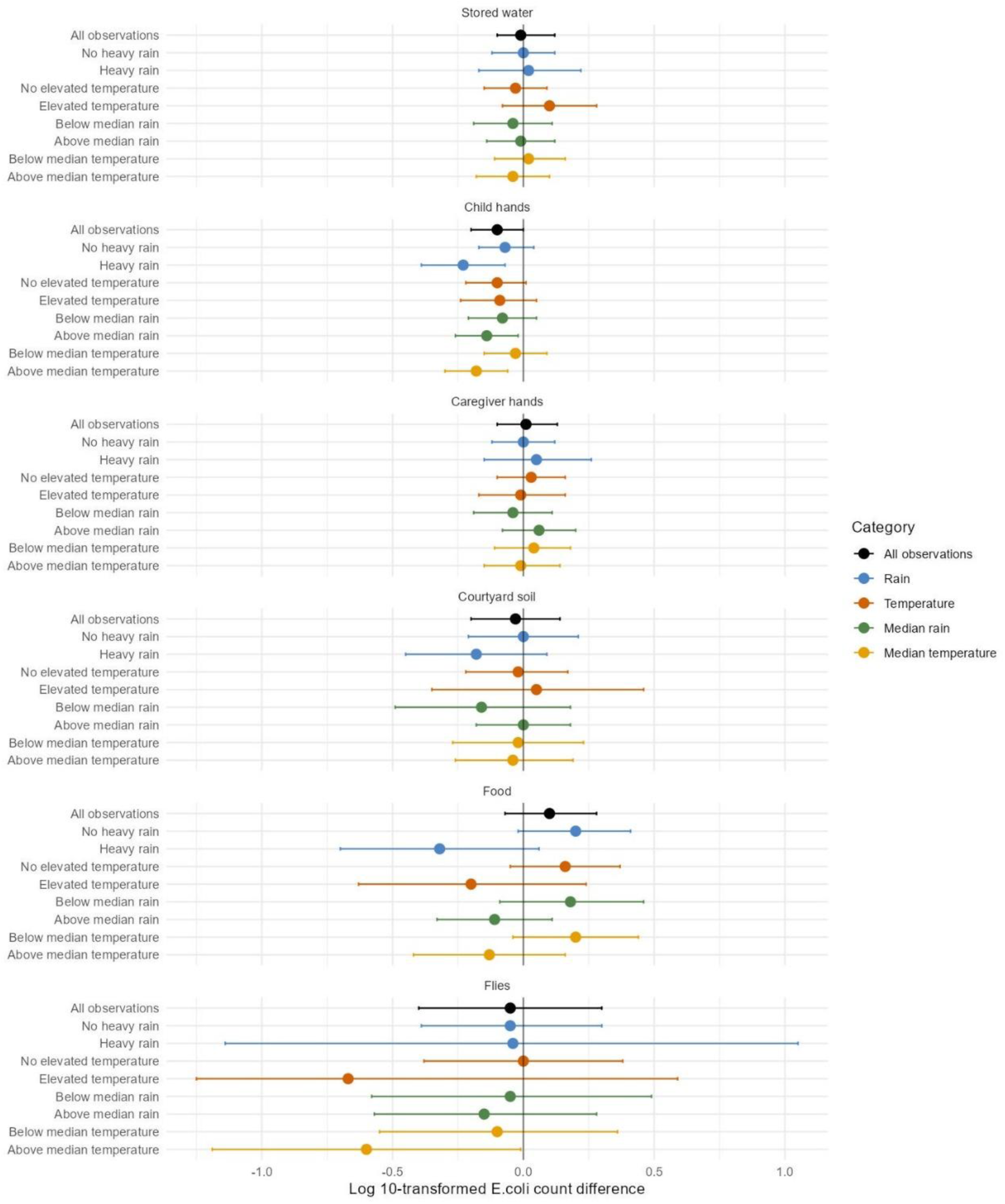
Effect modification by heavy rain, elevated temperature, above-vs. below-median rain, and above-vs. below-median temperature within 2 days before sampling on adjusted associations between finished floors and log 10-transformed *E. coli* counts in environmental samples. Heavy rain and elevated temperature are each defined as >80th percentile of daily values during the study period. Above-median rain and above-median temperature are each defined as >50th percentile of 2-day rolling-average rainfall and temperature values during the study period. The circles denote differences in log10-transformed *E. coli* counts between households with finished vs. unfinished floors, and the horizontal lines denote 95% confidence intervals. Confidence intervals for fly samples are truncated in the “elevated temperature” stratum because of low precision due to small sample size in this stratum. The numerical estimates corresponding to this figure are provided in **Table S6**.

Following heavy rain and above-median temperature, finished floors were associated with approximately 10% lower *E. coli* prevalence in stored drinking water (interaction p-value<0.20, Fig S3, Table S7). Finished floors appeared to be associated with lower *E. coli* prevalence in/on flies following periods of no heavy rain and below-median rain (interaction p-values <0.20, Fig S3, Table S7) but associations could not be distinguished from chance in any stratum. Finished floors were also associated with approximately 50% lower *E. coli* prevalence in/on flies following periods of elevated temperature (Fig S3, Table S7). There was limited evidence of effect modification by rainfall or temperature on *E. coli* prevalence in other sample types (Fig S3, Table S7).

#### Effect modification by animal ownership

In the bottom tertile of animal ownership, finished floors were associated with approximately 0.20 log higher *E. coli* counts on stored water and caregiver hands, and there was a similar trend for food that could not be distinguished from chance (Fig 4, Table S8). In the middle and top tertiles of animal ownership, finished floors were associated with approximately 0.20 log lower *E. coli* counts on child hands (interaction p-values <0.20 compared to the bottom tertile) (Fig 4, Table S8). Similarly, finished floors were associated with approximately 20% lower *E. coli* prevalence in child hand rinses compared to unfinished floors in the top tertile of animal ownership (interaction p-value<0.20 compared to bottom tertile, Fig S4, Table S9). There was insufficient evidence of effect modification by animal ownership categories on *E. coli* prevalence in other sample types (Fig S4, Table S9).

**Figure 4:**
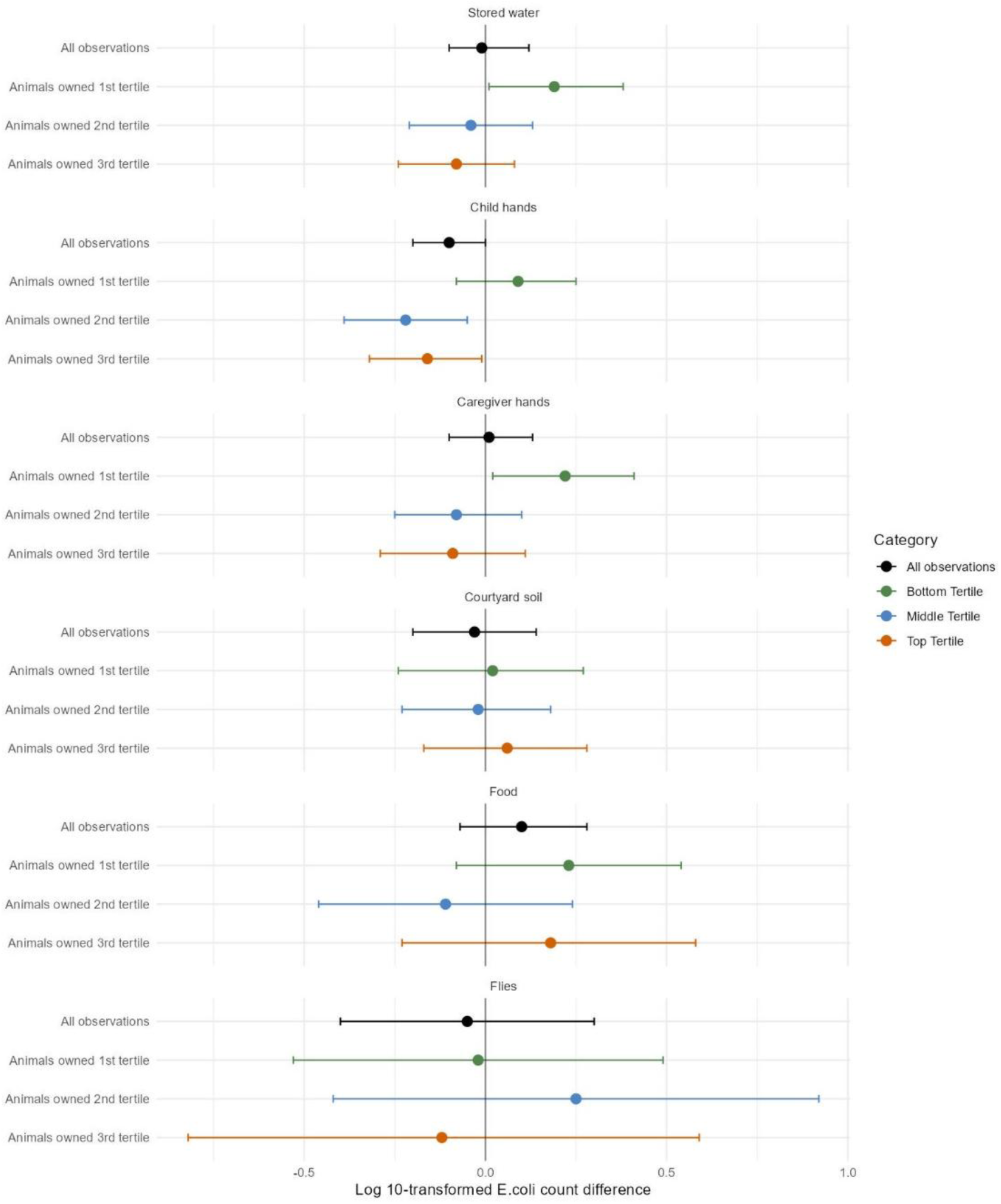
Effect modification across tertiles of animals owned on adjusted associations between finished floors and log 10-transformed *E. coli* counts in environmental samples. Households owned the following number of animals, mean (range): 1st tertile: 4.6 (0-9), 2nd tertile: 14.4 (10-19), 3rd tertile: 57.6 (20-2700). The circles denote differences in log10-transformed *E. coli* counts between households with finished vs. unfinished floors, and the horizontal lines denote 95% confidence intervals. The numerical estimates corresponding to this figure are provided in **Table S8**.

## Discussion

Among rural Bangladeshi households, having finished floors was associated with slightly lower log10-transformed *E. coli* counts (Δlog10 = -0.10 (-0.20, 0.00)) and prevalence (PR= 0.90 (0.83, 0.98)) on child hands compared to unfinished floors, adjusting for potential confounders (socio-demographics, water/sanitation status, animal ownership). Floor material was not associated with *E. coli* counts or prevalence in any other sample type after adjusting for potential confounders. Following periods of increased rainfall and temperature, finished floors were associated with up to 0.23-log lower *E. coli* counts on child hands, up to 0.33-log lower *E. coli* counts in food (but associations were not distinguishable from chance) and approximately 10% lower *E. coli* prevalence in stored drinking water. Finished floors also appeared to be associated with lower *E. coli* counts and prevalence on flies following periods of less rain and increased temperature.

A growing body of observational literature reports child health benefits associated with improved floor materials in low-income countries. In a matched cohort study in Mexico, replacing soil floors with concrete floors reduced diarrhea and intestinal parasite infections and improved cognitive development, suggesting that finished floors have the potential to improve acute as well as downstream outcomes.^23^ Even when households’ water or sanitation infrastructure was inadequate, finished floors were associated with reduced infant diarrheal illness in Zimbabwe compared to soil floors.^24^ Among children enrolled in the WASH Benefits trial in Kenya and Bangladesh, concrete floors were associated with reduced diarrhea and parasite infections caused by STH and Giardia.^39^ Our findings indicate that these health benefits in the WASH Benefits Bangladesh trial are not mediated by lower fecal contamination of the domestic environment as measured by *E. coli,* except for child hands. In Bangladesh, children’s mouthing of their own hands was the largest contributor to *E. coli* ingestion by children aged 6-35 months old.^4^ A recent meta-analysis found that the odds of child diarrhea increased by approximately 10% for each 1-log increase in *E. coli* on child hands,^45^ while in Bangladesh the incidence of child diarrhea increased by 23% for each 1-log increase in *E. coli* on child hands.^46^ Therefore, we expect lower *E. coli* contamination of children’s hands in homes with finished floors to result in reduced diarrheal illness. However, the associations we observed between floor material and *E. coli* on child hands were small in magnitude and unlikely to fully explain the previously reported protective associations between finished floors and child health outcomes in the WASH Benefits study.^46^

Following periods of increased rainfall and temperature, finished floors were associated with larger reductions in fecal contamination of child hands. There was also some evidence of larger reductions in contamination of food and stored drinking water but these associations should be interpreted with caution given the large number of subgroup analyses.^47, 48^ We have previously reported similar effect modification by rainfall and temperature for effects of WASH interventions on child health and fecal contamination in the WASH Benefits trial.^49, 50^ Bangladesh experiences a pronounced monsoon season, and heavy rainfall and/or flooding events are common. Our findings indicate that finished floors can reduce exposure to fecal organisms via child hands, food and stored water during these events. Rainfall and temperature extremes are expected to increase in the future,^51^ and finished floors may yield increased health benefits under climate change.

Finished floors also appeared to be associated with larger reductions in *E. coli* contamination of child hands but not caregiver hands in households that kept more vs. fewer domestic animals. While cohabitation with domestic animals is associated with higher levels of *E. coli* on floors regardless of floor material, our previous work indicates that, among households that keep animals, concrete floors harbor less *E. coli* than soil floors.^52^ Children have frequent hand contact with floors, and our current results indicate that improved floor materials may translate to reduced child hand contact with animal fecal contamination via floors. We did not observe a similar trend for caregiver hands. Women in rural Bangladesh often bear the primary responsibility for the daily upkeep and welfare of domestic animals^53^ and may have frequent direct hand contact with animal fecal waste; therefore, improved floor materials may not be sufficient to reduce *E. coli* contamination of caregiver hands.

Notably, all studies of health outcomes associated with floor material to date are observational, and confounding by socioeconomic status could have influenced their findings. In our analysis, households with finished floors were better educated, wealthier and more food secure than households with unfinished floors. While households with finished floors appeared to have lower *E. coli* contamination of stored drinking water, and caregiver and child hand rinses in unadjusted analyses, after adjusting for potential confounders, finished floors were only associated with lower *E. coli* counts and prevalence on child hands. While our analysis controlled for several sociodemographic indicators, it is possible that residual confounding remains. Future efforts to investigate the effects of finished floors on child health and environmental contamination should consider randomized study designs to minimize confounding.^54^

A strength of this study was the large sample size, with >20,000 samples collected across several fecal-oral transmission pathways over 3.5 years. The large sample size indicates that lack of observed associations is unlikely to be due to lack of statistical precision. However, some sample types (e.g., flies) were collected in smaller numbers than others and only approximately 10% of study households had finished floors, leading to small cells, especially in the effect modification analyses. Another weakness is that we relied on *E. coli* to measure fecal contamination. *E. coli* can originate from natural sources and does not correlate well with the presence of actual pathogens.^55^ Therefore, it is possible that the lack of associations we observed between floor material and *E. coli* do not apply to pathogens, and especially non-bacterial pathogens.

Additionally, while we investigated several fecal-oral transmission pathways, we did not measure *E. coli* directly on floors. It is possible that the health benefits associated with finished floors in previous studies were mediated by reduced direct exposure to pathogens on floors. Prior studies have demonstrated lower *E. coli* contamination on finished vs. unfinished floors. A study of peri-urban households in Peru enumerated *E. coli* on floors at the household entrance and kitchen areas.^17^ In both locations, dirt/soil floors had higher mean *E. coli* counts compared to wood or cement floors. Other studies in Bangladesh have detected fecal indicator bacteria such as *E. coli* on 25% of floors and enterococcus on 100% of floors, animal-specific fecal markers (BacR) on 27% of floors^56^ and antimicrobial-resistant *E. coli* on 71% of floors^39^ but these studies did not compare contamination between finished and unfinished floors. A study in Ghana found that floor material (concrete vs. soil) in the bedroom did not influence the prevalence and frequency of soil ingestion and the amount of soil ingested by children.^57^ We also note that even in houses with concrete floors, children can still be frequently exposed to soil and dust when they spend time outdoors (e.g. in courtyards).

In recent large trials, WASH interventions achieved limited child health improvements^37, 38^ and reductions in environmental contamination with pathogens.^58^ Finished floor materials hold promise as a potential intervention to improve child health in low-income countries. Concrete floors are easier to keep clean than soil floors. In contrast to WASH interventions, concrete floors, once installed, do not necessitate ongoing behavior change and are an aspirational product, which can potentially facilitate implementation by policymakers and NGOs. Our findings from rural Bangladesh indicate that finished floors may reduce exposure to fecal organisms via reduced contamination of child hands, and following periods of increased rainfall and temperature, via reduced contamination of child hands, food and stored drinking water. Measures to control enteric infections in low-income countries should test flooring improvements to reduce exposure to fecal contamination. Future studies should enumerate pathogens on finished vs. unfinished floors and use randomized study designs to eliminate the possibility of unmeasured confounding.

## Acknowledgement

This study was funded by the Bill & Melinda Gates Foundation (grant # OPPGD759), National Institutes of Health (grant # R01HD078912), and the World Bank. The authors declare no competing financial interest.

## Notes

### Competing Interest Statement

The authors have declared no competing interest.

## References

1. Kwong LH, Ercumen A, Pickering AJ, Unicomb L, Davis J, Luby SP. Hand- and Object-Mouthing of Rural Bangladeshi Children 3-18 Months Old. Int J Environ Res Public Health. 2016;13(6):563. doi:10.3390/ijerph13060563

2. Ngure FM, Humphrey JH, Mbuya MNN, et al. Formative research on hygiene behaviors and geophagy among infants and young children and implications of exposure to fecal bacteria. Am J Trop Med Hyg. 2013;89(4):709–716. doi:10.4269/ajtmh.12-0568

3. Kwong LH, Ercumen A, Pickering AJ, et al. Ingestion of Fecal Bacteria along Multiple Pathways by Young Children in Rural Bangladesh Participating in a Cluster-Randomized Trial of Water, Sanitation, and Hygiene Interventions (WASH Benefits). Environ Sci Technol. 2020;54(21):13828–13838. doi:10.1021/acs.est.0c02606

4. Morita T, Perin J, Oldja L, et al. Mouthing of Soil Contaminated Objects is Associated with Environmental Enteropathy in Young Children. Tropical Medicine & International Health. 2017;22(6):670–678. doi:10.1111/tmi.12869

5. George CM, Oldja L, Biswas S, et al. Geophagy is associated with environmental enteropathy and stunting in children in rural Bangladesh. Am J Trop Med Hyg. 2015;92(6):1117–1124. doi:10.4269/ajtmh.14-0672

6. Shivoga WA, Moturi WN. Geophagia as a risk factor for diarrhoea. J Infect Dev Ctries. 2009;3(02):094–098. doi:10.3855/jidc.55

7. Bauza V, Byrne DM, Trimmer JT, Lardizabal A, Atiim P, Asigbee MA, Guest JS. Child soil ingestion in rural Ghana–frequency, caregiver perceptions, relationship with household floor material and associations with child diarrhoea. Tropical Medicine & International Health. 2018 May;23(5):558–69.

8. Bauza V, Ocharo RM, Nguyen TH, Guest JS. Soil ingestion is associated with child diarrhea in an urban slum of Nairobi, Kenya. The American journal of tropical medicine and hygiene. 2017 Mar 3;96(3):569.

9. Colston JM, Fang B, Nong MK, et al. Spatial variation in housing construction material in low- and middle-income countries: a Bayesian spatial prediction model of a key infectious diseases risk factor and social determinant of health. medRxiv. Published online January 1, 2024:2024.05.23.24307833. doi:10.1101/2024.05.23.24307833

10. Halliday KE, Oswald WE, Mcharo C, et al. Community-level epidemiology of soil-transmitted helminths in the context of school-based deworming: Baseline results of a cluster randomised trial on the coast of Kenya. Garba A, ed. PLoS Negl Trop Dis. 2019;13(8):e0007427. doi:10.1371/journal.pntd.0007427

11. Krause RJ, Koski KG, Pons E, Sandoval N, Sinisterra O, Scott ME. Ascaris and hookworm transmission in preschool children from rural Panama: role of yard environment, soil eggs/larvae and hygiene and play behaviours. Parasitology. 2015;142(12):1543–1554. doi:10.1017/S0031182015001043

12. Basualdo JA, Córdoba MA, de Luca MM, et al. Intestinal parasitoses and environmental factors in a rural population of Argentina, 2002-2003. Rev Inst Med Trop Sao Paulo. 2007;49(4):251–255. doi:10.1590/s0036-46652007000400011

13. Benjamin-Chung J, Nazneen A, Halder AK, et al. The Interaction of Deworming, Improved Sanitation, and Household Flooring with Soil-Transmitted Helminth Infection in Rural Bangladesh. Lv S, ed. PLoS Negl Trop Dis. 2015;9(12):e0004256. doi:10.1371/journal.pntd.0004256

14. Hall A, Conway DJ, Anwar KS, Rahman ML. Strongyloides stercoralis in an urban slum community in Bangladesh: factors independently associated with infection. Trans R Soc Trop Med Hyg. 1994;88(5):527–530. doi:10.1016/0035-9203(94)90146-5

15. Cociancic P, Torrusio SE, Zonta ML, Navone GT. Risk factors for intestinal parasitoses among children and youth of Buenos Aires, Argentina. One Health. 2020;9:100116. doi:10.1016/j.onehlt.2019.100116

16. Brooker S, Clements ACA, Bundy DAP. Global Epidemiology, Ecology and Control of Soil-Transmitted Helminth Infections. In: Advances in Parasitology. Vol 62. Elsevier; 2006:221-261. doi:10.1016/S0065-308X(05)62007-6

17. Exum NG, Olórtegui MP, Yori PP, et al. Floors and Toilets: Association of Floors and Sanitation Practices with Fecal Contamination in Peruvian Amazon Peri-Urban Households. Environ Sci Technol. 2016;50(14):7373–7381. doi:10.1021/acs.est.6b01283

18. Navab-Daneshmand T, Friedrich MND, Gächter M, et al. Escherichia coli Contamination across Multiple Environmental Compartments (Soil, Hands, Drinking Water, and Handwashing Water) in Urban Harare: Correlations and Risk Factors. Am J Trop Med Hyg. 2018;98(3):803–813. doi:10.4269/ajtmh.17-0521

19. Pickering AJ, Julian TR, Marks SJ, et al. Fecal contamination and diarrheal pathogens on surfaces and in soils among Tanzanian households with and without improved sanitation. Environ Sci Technol. 2012;46(11):5736–5743. doi:10.1021/es300022c

20. Montealegre MC, Roy S, Böni F, et al. Risk Factors for Detection, Survival, and Growth of Antibiotic-Resistant and Pathogenic Escherichia coli in Household Soils in Rural Bangladesh. Vieille C, ed. Appl Environ Microbiol. 2018;84(24):e01978–18. doi:10.1128/AEM.01978-18

21. Ercumen A, Pickering AJ, Kwong LH, et al. Animal Feces Contribute to Domestic Fecal Contamination: Evidence from E. coli Measured in Water, Hands, Food, Flies, and Soil in Bangladesh. Environ Sci Technol. 2017;51(15):8725–8734. doi:10.1021/acs.est.7b01710

22. Thomson H, Thomas S, Sellstrom E, Petticrew M. The health impacts of housing improvement: a systematic review of intervention studies from 1887 to 2007. Am J Public Health. 2009;99 Suppl 3(Suppl 3):S681–692. doi:10.2105/AJPH.2008.143909

23. Cattaneo MD, Galiani S, Gertler PJ, Martinez S, Titiunik R. Housing, Health, and Happiness. American Economic Journal: Economic Policy. 2009;1(1):75–105. doi:10.1257/pol.1.1.75

24. Koyuncu A, Kang Dufour MS, Watadzaushe C, et al. Household flooring associated with reduced infant diarrhoeal illness in Zimbabwe in households with and without WASH interventions. Trop Med Int Health. 2020;25(5):635–643. doi:10.1111/tmi.13385

25. Legge H, Pullan RL, Sartorius B. Improved household flooring is associated with lower odds of enteric and parasitic infections in low- and middle-income countries: A systematic review and meta-analysis. Mutheneni SR, ed. PLOS Glob Public Health. 2023;3(12):e0002631. doi:10.1371/journal.pgph.0002631

26. Luby SP, Rahman M, Arnold BF, et al. Effects of water quality, sanitation, handwashing, and nutritional interventions on diarrhoea and child growth in rural Bangladesh: a cluster randomised controlled trial. The Lancet Global Health. 2018;6(3):e302–e315. doi:10.1016/S2214-109X(17)30490-4

27. Lin A, Ercumen A, Benjamin-Chung J, et al. Effects of Water, Sanitation, Handwashing, and Nutritional Interventions on Child Enteric Protozoan Infections in Rural Bangladesh: A Cluster-Randomized Controlled Trial. Clin Infect Dis. 2018;67(10):1515–1522. doi:10.1093/cid/ciy320

28. Ercumen A, Benjamin-Chung J, Arnold BF, et al. Effects of water, sanitation, handwashing and nutritional interventions on soil-transmitted helminth infections in young children: A cluster-randomized controlled trial in rural Bangladesh. Nery SV, ed. PLoS Negl Trop Dis. 2019;13(5):e0007323. doi:10.1371/journal.pntd.0007323

29. Contreras JD, Islam M, Mertens A, et al. Evaluation of an on-site sanitation intervention against childhood diarrhea and acute respiratory infection 1 to 3.5 years after implementation: Extended follow-up of a cluster-randomized controlled trial in rural Bangladesh. PLoS Med. 2022;19(8):e1004041. doi:10.1371/journal.pmed.1004041

30. Ercumen A, Mertens A, Arnold BF, et al. Effects of Single and Combined Water, Sanitation and Handwashing Interventions on Fecal Contamination in the Domestic Environment: A Cluster-Randomized Controlled Trial in Rural Bangladesh. Environ Sci Technol. 2018;52(21):12078–12088. doi:10.1021/acs.est.8b05153

31. Ercumen A, Pickering AJ, Kwong LH, et al. Do Sanitation Improvements Reduce Fecal Contamination of Water, Hands, Food, Soil, and Flies? Evidence from a Cluster-Randomized Controlled Trial in Rural Bangladesh. Environ Sci Technol. 2018;52(21):12089–12097. doi:10.1021/acs.est.8b02988

32. Fuhrmeister ER, Ercumen A, Pickering AJ, et al. Effect of Sanitation Improvements on Pathogens and Microbial Source Tracking Markers in the Rural Bangladeshi Household Environment. Environ Sci Technol. 2020;54(7):4316–4326. doi:10.1021/acs.est.9b04835

33. Contreras JD, Islam M, Mertens A, et al. Longitudinal Effects of a Sanitation Intervention on Environmental Fecal Contamination in a Cluster-Randomized Controlled Trial in Rural Bangladesh. Environ Sci Technol. 2021;55(12):8169–8179. doi:10.1021/acs.est.1c01114

34. Pickering AJ, Njenga SM, Steinbaum L, et al. Effects of single and integrated water, sanitation, handwashing, and nutrition interventions on child soil-transmitted helminth and Giardia infections: A cluster-randomized controlled trial in rural Kenya. PLoS Med. 2019;16(6):e1002841. doi:10.1371/journal.pmed.1002841

35. Null C, Stewart CP, Pickering AJ, et al. Effects of water quality, sanitation, handwashing, and nutritional interventions on diarrhoea and child growth in rural Kenya: a cluster-randomised controlled trial. The Lancet Global Health. 2018;6(3):e316–e329. doi:10.1016/S2214-109X(18)30005-6

36. Swarthout JM, Mureithi M, Mboya J, et al. Addressing Fecal Contamination in Rural Kenyan Households: The Roles of Environmental Interventions and Animal Ownership. Environ Sci Technol. 2024;58(22):9500–9514. doi:10.1021/acs.est.3c09419

37. Pickering AJ, Null C, Winch PJ, et al. The WASH Benefits and SHINE trials: interpretation of WASH intervention effects on linear growth and diarrhoea. The Lancet Global Health. 2019;7(8):e1139–e1146. doi:10.1016/S2214-109X(19)30268-2

38. Cumming O, Arnold BF, Ban R, et al. The implications of three major new trials for the effect of water, sanitation and hygiene on childhood diarrhea and stunting: a consensus statement. BMC Medicine. 2019;17(1):173. doi:10.1186/s12916-019-1410-x

39. Benjamin-Chung J, Crider YS, Mertens A, et al. Household finished flooring and soil-transmitted helminth and Giardia infections among children in rural Bangladesh and Kenya: a prospective cohort study. The Lancet Global Health. 2021;9(3):e301–e308. doi:10.1016/S2214-109X(20)30523-4

40. Arnold BF, Null C, Luby SP, et al. Cluster-randomised controlled trials of individual and combined water, sanitation, hygiene and nutritional interventions in rural Bangladesh and Kenya: the WASH Benefits study design and rationale. BMJ Open. 2013;3(8):e003476. doi:10.1136/bmjopen-2013-003476

41. Yakub GP, Castric DA, Stadterman-Knauer KL, et al. Evaluation of Colilert and Enterolert Defined Substrate Methodology for Wastewater Applications. Water Environment Research. 2002;74(2):131–135. doi:10.2175/106143002X139839

42. Coates J, Swindale A, Bilinsky P. Household Food Insecurity Access Scale (HFIAS) for Measurement of Household Food Access: Indicator Guide Version 3. 2007. Food and Nutrition Technical Assistance Project Academy for Educational Development, Washington DC.

43. Mertens Andrew, Balakrishnan Kalpana, Ramaswamy Padmavathi, et al. Associations between High Temperature, Heavy Rainfall, and Diarrhea among Young Children in Rural Tamil Nadu, India: A Prospective Cohort Study. Environmental Health Perspectives. 127(4):047004. doi:10.1289/EHP3711

44. Niven C, Islam M, Nguyen A, et al. Effects of weather extremes on fecal contamination along pathogen transmission pathways in rural Bangladeshi households. Published online December 30, 2023. doi:10.1101/2023.12.27.23300582

45. Goddard FGB, Pickering AJ, Ercumen A, Brown J, Chang HH, Clasen T. Faecal contamination of the environment and child health: a systematic review and individual participant data meta-analysis. The Lancet Planetary Health. 2020;4(9):e405–e415. doi:10.1016/S2542-5196(20)30195-9

46. Pickering AJ, Ercumen A, Arnold BF, et al. Fecal Indicator Bacteria along Multiple Environmental Transmission Pathways (Water, Hands, Food, Soil, Flies) and Subsequent Child Diarrhea in Rural Bangladesh. Environ Sci Technol. 2018;52(14):7928–7936. doi:10.1021/acs.est.8b00928

47. Lagakos SW. The Challenge of Subgroup Analyses — Reporting without Distorting. N Engl J Med. 2006;354(16):1667–1669. doi:10.1056/NEJMp068070

48. Burke JF, Sussman JB, Kent DM, Hayward RA. Three simple rules to ensure reasonably credible subgroup analyses. BMJ. Published online November 4, 2015:h5651. doi:10.1136/bmj.h5651

49. Nguyen AT, Grembi JA, Riviere M, Barratt Heitmann G, Hutson WD, Athni TS, Patil A, Ercumen A, Lin A, Crider Y, Mertens A. Influence of temperature and precipitation on the effectiveness of water, sanitation, and handwashing interventions against childhood diarrheal disease in rural Bangladesh: a reanalysis of the WASH Benefits Bangladesh trial. Environmental Health Perspectives. 2024 Apr 11;132(4):047006.

50. Niven CG, Islam M, Nguyen A, Mertens A, Pickering AJ, Kwong LH, Alam M, Sen D, Islam S, Rahman M, Unicomb L. Rainfall and Temperature Modify Effects of On-Site Sanitation Intervention on E. coli Contamination in Bangladeshi Households. medRxiv. 2024:2024–08.

51. Myhre G, Alterskjær K, Stjern CW, et al. Frequency of extreme precipitation increases extensively with event rareness under global warming. Sci Rep. 2019;9(1):16063. doi:10.1038/s41598-019-52277-4

52. Ercumen A, Hossain MdS, Tabassum T, et al. Dirt floors and domestic animals are associated with soilborne exposure to antimicrobial resistant E. coli in rural Bangladeshi households. Published online February 23, 2025. doi:10.1101/2025.02.21.639507

53. Mostari MP, Sadrul SB, Rahman MH, Islam MS. Women Empowerment and Livestock Development in Bangladesh: A Review. Bangladesh J of Livestock Res. Published online December 1, 2021:1–15. doi:10.3329/bjlr.v28i1.72014

54. Rahman M, Jahan F, Hanif S, et al. Effects of household concrete floors on maternal and child health – the CRADLE trial: a randomised controlled trial protocol. Published online July 27, 2024. doi:10.1101/2024.07.26.24311076

55. Wu J, Long SC, Das D, Dorner SM. Are microbial indicators and pathogens correlated? A statistical analysis of 40 years of research. Journal of water and health. 2011 Jun 1;9(2):265–78.

56. Harris AR, Pickering AJ, Harris M, Doza S, Islam MS, Unicomb L, Luby S, Davis J, Boehm AB. Ruminants contribute fecal contamination to the urban household environment in Dhaka, Bangladesh. Environmental science & technology. 2016 May 3;50(9):4642–9.

57. Tabassum T, Hossain MdS, Ercumen A, et al. Isolation and characterization of cefotaxime resistant Escherichia coli from household floors in rural Bangladesh. Heliyon. 2024;10(14). doi:10.1016/j.heliyon.2024.e34367

58. Mertens A, Arnold BF, Benjamin-Chung J, et al. Effects of water, sanitation, and hygiene interventions on detection of enteropathogens and host-specific faecal markers in the environment: a systematic review and individual participant data meta-analysis. The Lancet Planetary Health. 2023;7(3):e197–e208. doi:10.1016/S2542-5196(23)00028-1

