## supporting information for "Associations between floor material and *E. coli* contamination in rural Bangladeshi households"

[**Figure S2:** Associations between floor material and prevalence of *E. coli* environmental samples by sample type. Unadjusted and adjusted prevalence ratios (PRs) for finished vs. unfinished floors, 95% confidence intervals (CIs) and associated p-values are shown for each sample. Adjusted analyses controlled for study arm (intervention vs. control), sex and age of index child, caregiver’s age and education, number of children <18 years in the household, number of individuals living in the compound, food security, asset-based wealth index, wall materials, drinking water source, minutes to the primary drinking water source, number of cows, chickens and sheep/goats, narrow mouth container (only for stored water samples), cover status of the container (only for stored water samples), hours since water has been stored (only for stored water samples), and hours since stored food was prepared (only for food samples).	5](#_Toc194664088)

[**Figure S3:** Effect modification by heavy rain, elevated temperature, above- vs. below-median rain, and above- vs. below-median temperature within 2 days before sampling on adjusted associations between finished floors and *E. coli* prevalence in environmental samples. Heavy rain and elevated temperature are each defined as >80th percentile of daily values during the study period. Above-median rain and above-median temperature are each defined as >50th percentile of 2-day rolling-average rainfall and temperature values during the study period. The circles denote prevalence ratios for households with finished vs. unfinished floors and the horizontal lines denote 95% confidence intervals. The numerical estimates corresponding to this figure are provided in **Table S7.**	6](#_Toc194664089)

[**Table S6.** Effect modification by rainfall and temperature within 2 days prior to sampling on adjusted associations between flooring material and log 10-transformed *E. coli* counts in environmental samples. Heavy rain and elevated temperature are each defined as > 80th percentile of daily values during the study period. Above-median rain and temperature are defined as rolling 2-day average values >50^th^ percentile. Interaction p-values <0.20 are denoted in bold and interpreted as evidence of effect modification. The estimates in this table correspond to **Figure 3** in the main text.	14](#_Toc194664096)

[**Table S7.** Effect modification by rainfall and temperature within 2 days prior to sampling on adjusted associations between floor material and *E. coli* prevalence in environmental samples. Heavy rain and elevated temperature are each defined as > 80th percentile of daily values during the study period. Above-median rain and temperature are defined as rolling 2-day average values >50^th^ percentile. Interaction p-values <0.20 are denoted in bold and interpreted as evidence of effect modification. The estimates in this table correspond to **Figure S3**.	16](#_Toc194664097)

[**Table S8.** Effect modification by tertiles of animals owned on adjusted associations between flooring material and log 10-transformed *E. coli* counts in environmental samples. Households owned the following number of animals, mean (range): 1st tertile: 4.56 (0-9), 2nd tertile: 14.38 (10-19), 3rd tertile: 57.62 (20-2700). Interaction p-values <0.20 are denoted in bold and interpreted as evidence of effect modification. The estimates in this table correspond to **Figure 4** in the main text.	18](#_Toc194664098)

[**Table S9.** Effect modification by tertiles of animals owned on adjusted associations between floor material and *E. coli* prevalence in environmental samples. Households owned the following number of animals, mean (range): 1st tertile: 4.56 (0-9), 2^nd^ tertile: 14.38 (10-19), 3^rd^ tertile: 57.62 (20-2700). Interaction p-values <0.20 are denoted in bold and interpreted as evidence of effect modification. The estimates in this table correspond to **Figure S4**.	19](#_Toc194664099)

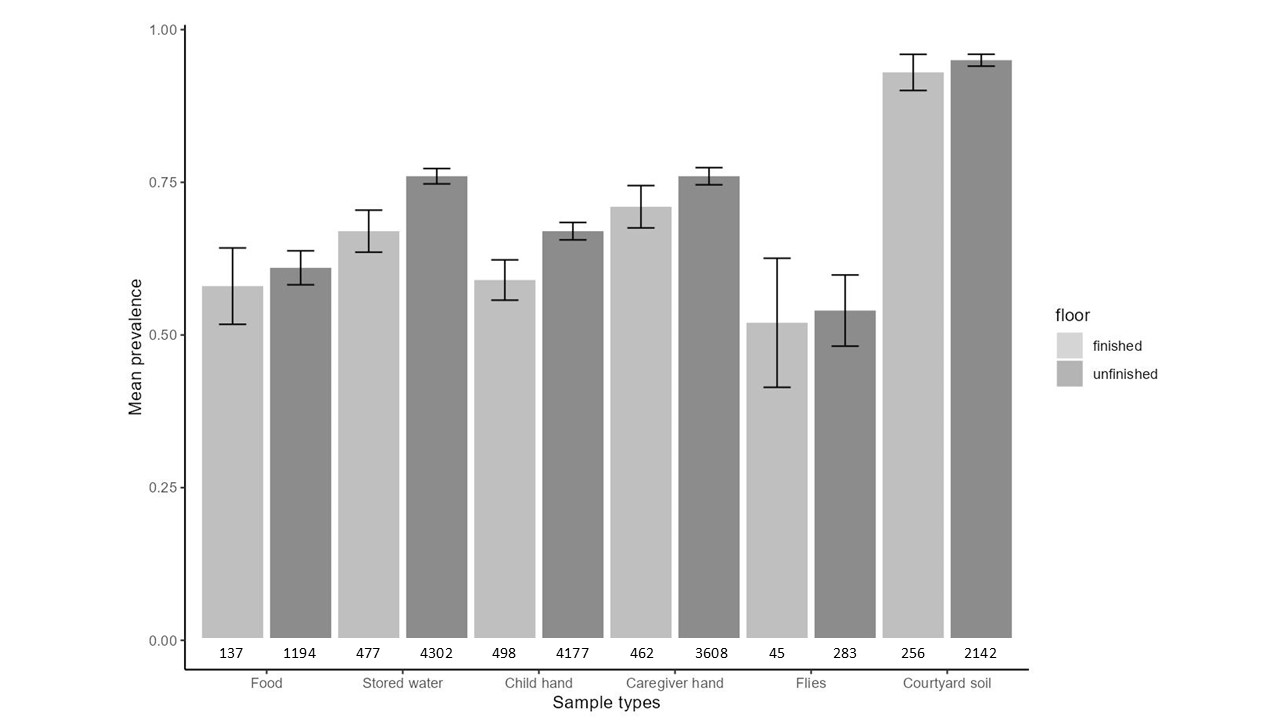

### **Figure S1**: Prevalence of *E. coli* with 95% confidence intervals (CI) by sample type and floor material. Each pair of columns represents a sample type. The number of samples is listed below the columns. Units by sample type are per one dry gram of food, 100 mL of stored water, two child/caregiver hands, one fly and one dry gram of soil.

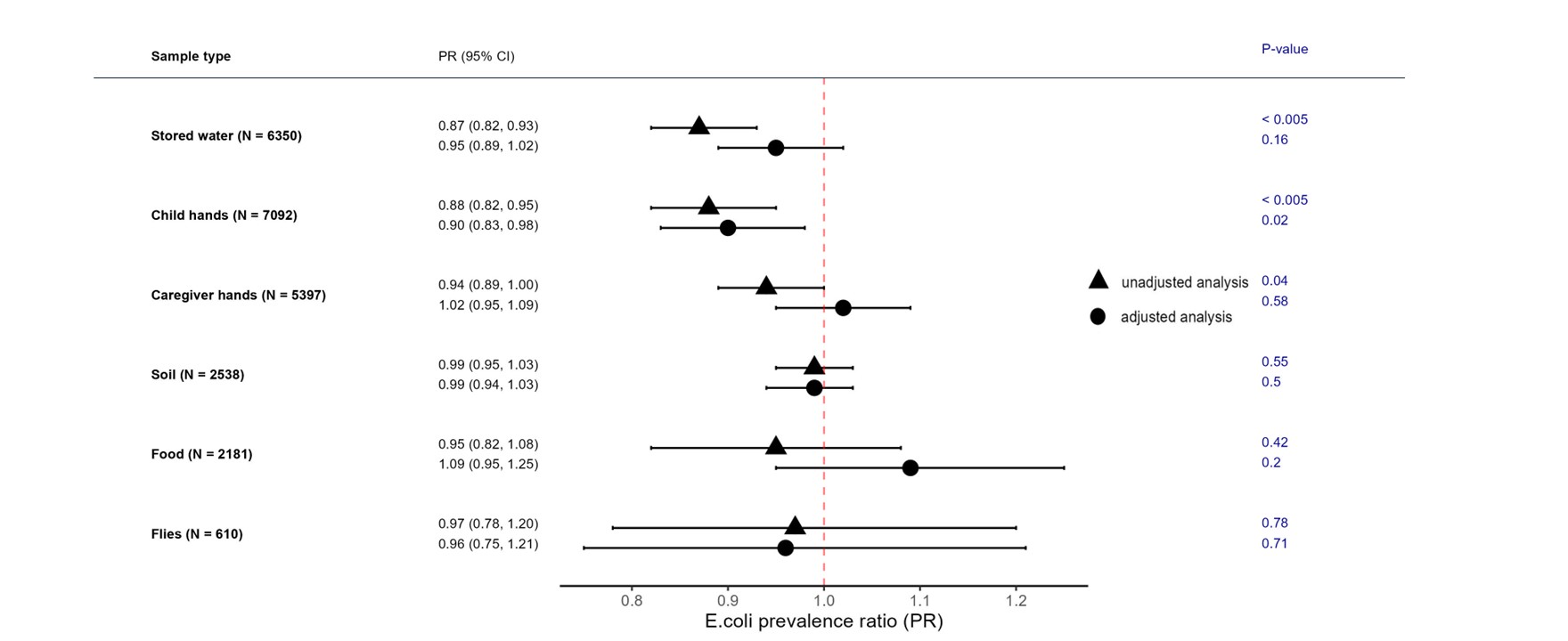

### **Figure S2:** Associations between floor material and prevalence of *E. coli* environmental samples by sample type. Unadjusted and adjusted prevalence ratios (PRs) for finished vs. unfinished floors, 95% confidence intervals (CIs) and associated p-values are shown for each sample. Adjusted analyses controlled for study arm (intervention vs. control), sex and age of index child, caregiver’s age and education, number of children <18 years in the household, number of individuals living in the compound, food security, asset-based wealth index, wall materials, drinking water source, minutes to the primary drinking water source, number of cows, chickens and sheep/goats, narrow mouth container (only for stored water samples), cover status of the container (only for stored water samples), hours since water has been stored (only for stored water samples), and hours since stored food was prepared (only for food samples).

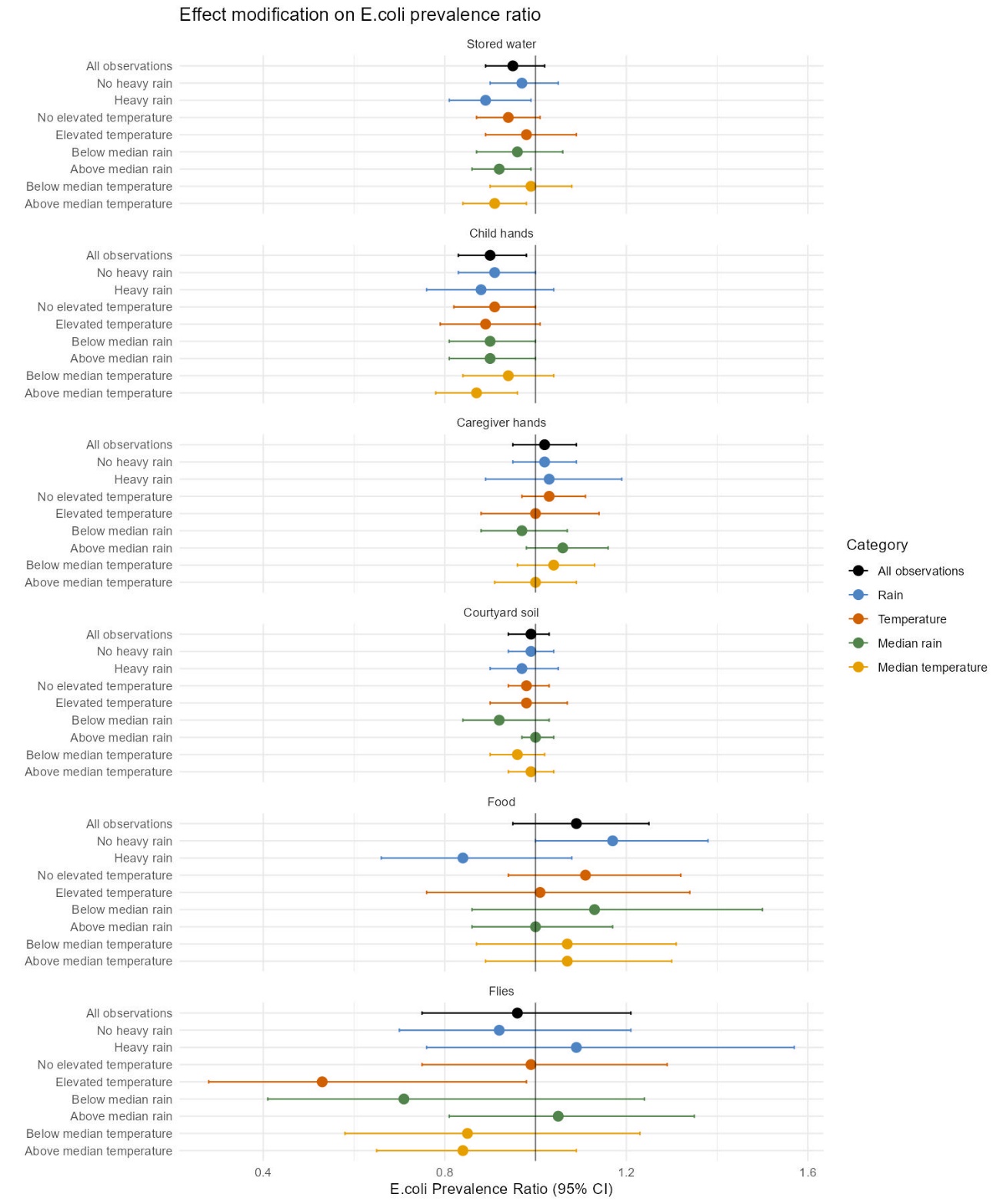

### **Figure S3:** Effect modification by heavy rain, elevated temperature, above- vs. below-median rain, and above- vs. below-median temperature within 2 days before sampling on adjusted associations between finished floors and *E. coli* prevalence in environmental samples. Heavy rain and elevated temperature are each defined as >80th percentile of daily values during the study period. Above-median rain and above-median temperature are each defined as >50th percentile of 2-day rolling-average rainfall and temperature values during the study period. The circles denote prevalence ratios for households with finished vs. unfinished floors and the horizontal lines denote 95% confidence intervals. The numerical estimates corresponding to this figure are provided in **Table S7.**

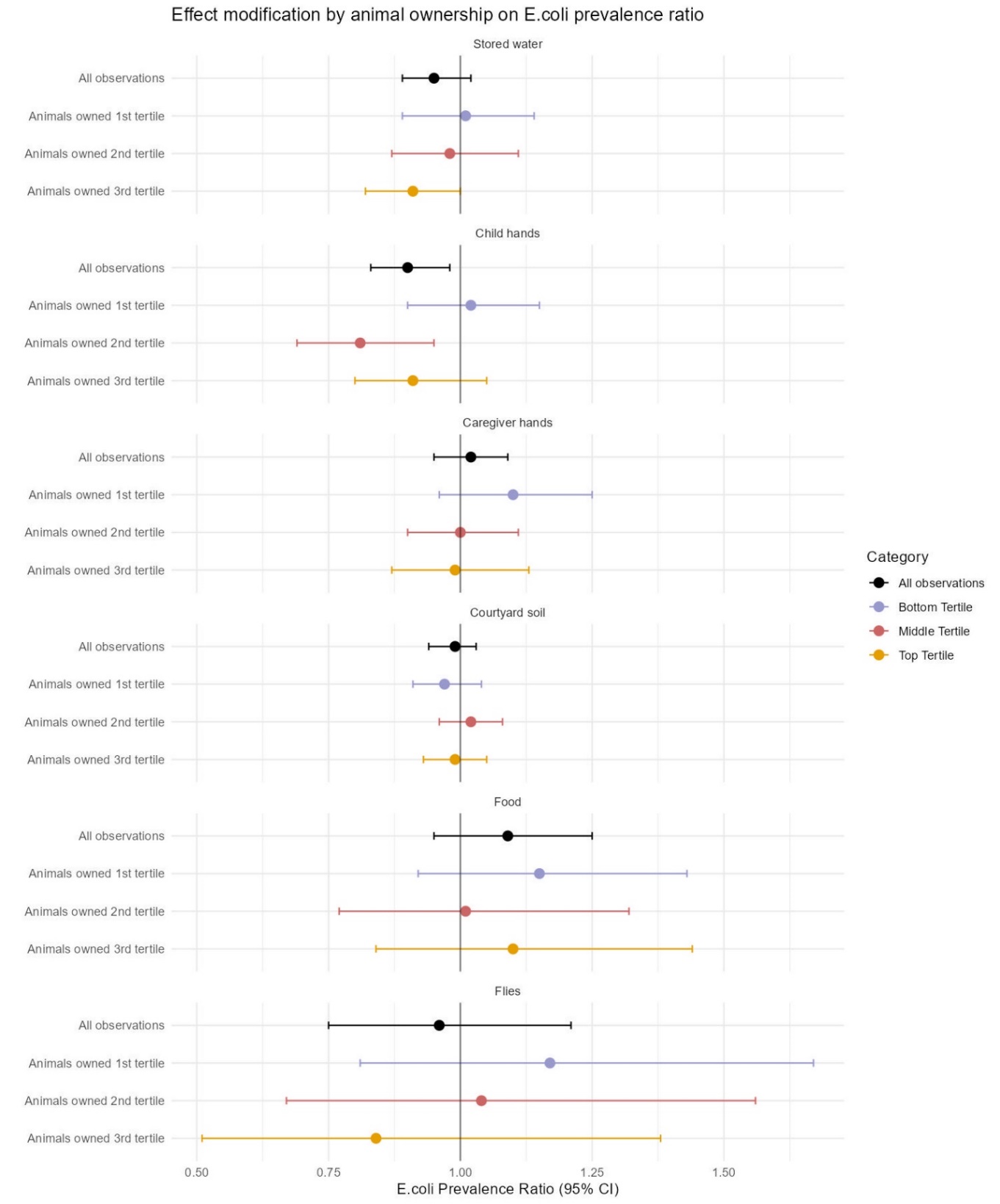

### **Figure S4:** Effect modification across tertiles of animals owned on adjusted associations between finished floors and *E. coli* prevalence in environmental samples. Households owned the following number of animals, mean (range): 1st tertile: 4.6 (0-9), 2^nd^ tertile: 14.4 (10-19), 3^rd^ tertile: 57.6 (20-2700). The circles denote prevalence ratios for households with finished vs. unfinished floors and the horizontal lines denote 95% confidence intervals. The numerical estimates corresponding to this figure are provided in **Table S9**.

### **Table S1.** Summary of household participation and sample collection over nine sampling rounds

| **Rounds** | **Number of household visits** | | **Number of samples collected by sample type** | | | | | | | | | | | |
| --- | --- | --- | --- | --- | --- | --- | --- | --- | --- | --- | --- | --- | --- | --- |
|  |  |  | **Stored water** | | **Child hands** | | **Caregiver hands** | | **Courtyard soil** | | **Food** | | **Flies** | |
|  | **F** | **U** | **F** | **U** | **F** | **U** | **F** | **U** | **F** | **U** | **F** | **U** | **F** | **U** |
| 1 | 212 | 1615 | 184 | 1439 | 210 | 1558 | – | – | 199 | 1596 | 186 | 1460 | 68 | 754 |
| 2 | 86 | 634 | 71 | 570 | 86 | 634 | 86 | 634 | – | – | – | – | – | – |
| 3 | 84 | 621 | 69 | 523 | 84 | 621 | 84 | 621 | – | – | – | – | – | – |
| 4 | 83 | 604 | 62 | 509 | 82 | 600 | 81 | 603 | 41 | 361 | 31 | 298 | – | – |
| 5 | 83 | 601 | 73 | 526 | 82 | 591 | 82 | 600 | 34 | 307 | 19 | 187 | – | – |
| 6 | 81 | 591 | 62 | 519 | 78 | 575 | 80 | 588 | – | – | – | – | – | – |
| 7 | 81 | 587 | 66 | 516 | 80 | 570 | 80 | 582 | – | – | – | – | – | – |
| 8 | 79 | 572 | 65 | 509 | 72 | 554 | 76 | 567 | – | – | – | – | – | – |
| 9 | 79 | 560 | 65 | 522 | 73 | 542 | 78 | 555 | – | – | – | – | – | – |
| **Total** | **868** | **6385** | **717** | **5633** | **847** | **6245** | **647** | **4750** | **274** | **2264** | **236** | **1945** | **86** | **524** |

F: households with finished floor, U: households with unfinished floor.

### **Table S2**. Differences in log 10-transformed *E. coli* counts in environmental samples for households with finished vs. unfinished floors. The estimates in this table correspond to **Figure 2** in the main text.

| **Sample type** | **N** | **Unfinished floors** | | **Finished floors** | | **Unadjusted analysis** | | **Adjusted analysis ^a^** | |
| --- | --- | --- | --- | --- | --- | --- | --- | --- | --- |
|  |  | **N** | **Log10 Mean MPN (SD)** | **N** | **Log10 Mean MPN (SD)** | **Log10 Difference (95% CI) ^b^** | **p-value** | **Log10 Difference (95% CI) ^b^** | **p-value** |
| Stored water | 6350 | 5633 | 0.92 (1.05) | 717 | 0.71 (1.05) | **-0.21 (-0.31, -0.11)** | **<0.005** | -0.01 (-0.10, 0.12) | 0.90 |
| Child hands | 7092 | 6245 | 1.33 (0.99) | 847 | 1.16 (0.92) | **-0.17 (-0.26, -0.09)** | **<0.005** | **-0.10 (-0.20, 0.00)** | **0.05** |
| Caregiver hands | 5397 | 4750 | 1.49 (1.01) | 647 | 1.35 (0.93) | **-0.13 (-0.23, -0.04)** | **<0.005** | 0.01 (-0.10, 0.13) | 0.85 |
| Courtyard soil | 2538 | 2264 | 5.11 (1.06) | 274 | 4.96 (1.08) | -0.15 (-0.32, 0.02) | 0.08 | -0.03 (-0.20, 0.14) | 0.72 |
| Food | 2181 | 1945 | 0.70 (1.40) | 236 | 0.63 (1.38) | -0.07 (-0.26, 0.12) | 0.45 | 0.10 (-0.07, 0.28) | 0.25 |
| Flies | 610 | 524 | 2.85 (1.35) | 86 | 2.85 (1.34) | -0.01 (-0.35, 0.33) | 0.96 | -0.05 (-0.40, 0.30) | 0.77 |

SD: standard deviation, CI: confidence interval.

^a^ Adjusted analyses controlled for study arm (intervention vs. control), sex and age of index child, caregiver’s age and education, number of children <18 years in the household, number of individuals living in the compound, food security, asset-based wealth index, wall materials, drinking water source, minutes to the primary drinking water source, number of cows, chickens and sheep/goats, narrow mouth container (only for stored water samples), cover status of the container (only for stored water samples), hours since water has been stored (only for stored water samples), and hours since stored food was prepared (only for food samples). Units by sample type are per 100 mL of stored water, two child/caregiver hands, one dry gram of soil/food, and one fly.

^b^ Estimates with p-values<0.05 are denoted in bold.

### **Table S3.** *E. coli* prevalence ratios in environmental samples for households with finished vs. unfinished floors. The estimates in this table correspond to **Figure S2**.

| **Sample type** | **N** | **Unfinished floors** | | **Finished floors** | | **Unadjusted analysis** | | **Adjusted analysis ^a^** | |
| --- | --- | --- | --- | --- | --- | --- | --- | --- | --- |
|  |  | **N** | **Prev % (n)** | **N** | **Prev % (n)** | **PR (95% CI) ^b^** | **p-value** | **PR (95% CI) ^b^** | **p-value** |
| Stored water | 6350 | 5633 | 76.4 (4302) | 717 | 66.5 (477) | **0.87 (0.82, 0.93)** | **<0.005** | 0.95 (0.89, 1.02) | 0.16 |
| Child hands | 7092 | 6245 | 66.9 (4177) | 847 | 58.8 (498) | **0.88 (0.82, 0.95)** | **<0.005** | **0.90 (0.83, 0.98)** | **0.02** |
| Caregiver hands | 5397 | 4750 | 76.0 (3608) | 647 | 71.4 (462) | **0.94 (0.89, 1.00)** | **0.04** | 1.02 (0.95, 1.09) | 0.58 |
| Courtyard soil | 2538 | 2264 | 94.6 (2142) | 274 | 93.4 (256) | 0.99 (0.95, 1.03) | 0.55 | 0.99 (0.94, 1.03) | 0.50 |
| Food | 2181 | 1945 | 61.4 (1194) | 236 | 58.1 (137) | 0.95 (0.82, 1.08) | 0.42 | 1.09 (0.95, 1.25) | 0.20 |
| Flies | 610 | 524 | 54.0 (283) | 86 | 52.3 (45) | 0.97 (0.78, 1.20) | 0.78 | 0.96 (0.75, 1.21) | 0.71 |

PR: prevalence ratio, CI: confidence interval.

^a^ Adjusted analyses controlled for study arm (intervention vs. control), sex and age of index child, caregiver’s age and education, number of children <18 years in the household, number of individuals living in the compound, food security, asset-based wealth index, wall materials, drinking water source, minutes to the primary drinking water source, number of cows, chickens and sheep/goats, narrow mouth container (only for stored water samples), cover status of the container (only for stored water samples), hours since water has been stored (only for stored water samples), and hours since stored food was prepared (only for food samples). Units by sample type are per 100 mL of stored water, two child/caregiver hands, one dry gram of soil/food, and one fly.

^b^ Estimates with p-values<0.05 are denoted in bold.

### **Table S4.** Sensitivity analysis for adjusted differences in log 10-transformed *E. coli* counts in environmental samples for households with finished vs. unfinished floors, using data from the subset of households which had the same floor status throughout the study.

| **Sample type** | **N** | **Primary analysis** | | **N** | **Sensitivity analysis** | |
| --- | --- | --- | --- | --- | --- | --- |
|  |  | **Log10 Difference (95% CI)** | **p-value** |  | **Log10 Difference (95% CI)** | **p-value** |
| Stored water | 6350 | -0.01 (-0.10, 0.12) | 0.90 | 5625 | 0.03 (-0.08, 0.15) | 0.54 |
| Child hands | 7092 | **-0.10 (-0.20, 0.00)** | **0.05** | 6324 | -0.07 (-0.18, 0.05) | 0.25 |
| Caregiver hands | 5397 | 0.01 (-0.10, 0.13) | 0.85 | 4817 | 0.01 (-0.13, 0.14) | 0.93 |
| Courtyard soil | 2538 | -0.03 (-0.20, 0.14) | 0.72 | 2205 | 0.00 (-0.19, 0.20) | 0.99 |
| Food | 2181 | 0.10 (-0.07, 0.28) | 0.25 | 1890 | 0.11 (-0.10, 0.33) | 0.30 |
| Flies | 610 | -0.05 (-0.40, 0.30) | 0.77 | 523 | -0.03 (-0.44, 0.38) | 0.89 |

CI: confidence interval

### **Table S5.** Sensitivity analysis for adjusted *E. coli* prevalence ratio in environmental samples for households with finished vs. unfinished floors, using data from the subset of households which had the same floor status throughout the study.

| **Sample type** | **N** | **Primary analysis** | | **N** | **Sensitivity analysis** | |
| --- | --- | --- | --- | --- | --- | --- |
|  |  | **PR (95% CI)** | **p-value** |  | **PR (95% CI)** | **p-value** |
| Stored water | 6350 | 0.95 (0.89, 1.02) | 0.16 | 5625 | 0.96 (0.90, 1.03) | 0.31 |
| Child hands | 7092 | **0.90 (0.83, 0.98)** | **0.02** | 6324 | 0.92 (0.83, 1.02) | 0.13 |
| Caregiver hands | 5397 | 1.02 (0.95, 1.09) | 0.58 | 4817 | 1.02 (0.94, 1.10) | 0.66 |
| Courtyard soil | 2538 | 0.99 (0.94, 1.03) | 0.50 | 2205 | 0.99 (0.94, 1.04) | 0.63 |
| Food | 2181 | 1.09 (0.95, 1.25) | 0.20 | 1890 | 1.10 (0.95, 1.27) | 0.21 |
| Flies | 610 | 0.96 (0.75, 1.21) | 0.71 | 523 | 0.99 (0.75, 1.30) | 0.93 |

PR: prevalence ratio, CI: confidence interval

### **Table S6.** Effect modification by rainfall and temperature within 2 days prior to sampling on adjusted associations between flooring material and log 10-transformed *E. coli* counts in environmental samples. Heavy rain and elevated temperature are each defined as > 80th percentile of daily values during the study period. Above-median rain and temperature are defined as rolling 2-day average values >50^th^ percentile. Interaction p-values <0.20 are denoted in bold and interpreted as evidence of effect modification. The estimates in this table correspond to **Figure 3** in the main text.

|  | **N** | | | **Log10 Mean MPN (SD)** | | **ΔLog10 (95% CI)** | **p-value** | **Interaction**  **p-value** |
| --- | --- | --- | --- | --- | --- | --- | --- | --- |
|  | **Total** | **Unfinished** | **Finished** | **Unfinished** | **Finished** |  |  |  |
| **Stored water** |  |  |  |  |  |  |  |  |
| All observations | 6350 | 5633 | 717 | 0.92 (1.05) | 0.71 (1.05) | -0.01 (-0.10, 0.12) | 0.90 | -- |
| No heavy rain | 5085 | 4527 | 558 | 0.85 (1.04) | 0.64 (1.01) | 0.00 (-0.12, 0.12) | 0.96 |  |
| Heavy rain | 1265 | 1106 | 159 | 1.18 (1.04) | 0.94 (1.17) | 0.02 ( -0.17, 0.22) | 0.81 | 0.70 |
| No elevated temperature | 4723 | 4217 | 506 | 0.87 (1.04) | 0.64 (1.04) | -0.03 (-0.15, 0.09) | 0.61 |  |
| Elevated temperature | 1627 | 1416 | 211 | 1.06 (1.07) | 0.86 (1.06) | 0.10 (-0.08, 0.28) | 0.29 | 0.72 |
| Below median rain | 3222 | 2906 | 316 | 0.72 (0.99) | 0.52 (0.94) | -0.04 (-0.19. 0.11) | 0.58 |  |
| Above median rain | 3128 | 2727 | 401 | 1.12 (1.08) | 0.85 (1.11) | -0.01 (-0.14, 0.12) | 0.94 | 0.63 |
| Below median temperature | 3488 | 3140 | 348 | 0.77 (1.01) | 0.64 (1.03) | 0.02 (-0.11, 0.16) | 0.74 |  |
| Above median temperature | 2862 | 2493 | 369 | 1.10 (1.07) | 0.77 (1.07) | -0.04 (-0.18, 0.10) | 0.55 | **0.02** |
| **Child hands** |  |  |  |  |  |  |  |  |
| All observations | 7092 | 6245 | 847 | 1.33 (0.99 | 1.16 (0.92) | **-0.10 (-0.20, 0.00)** | 0.05 | -- |
| No heavy rain | 5657 | 4996 | 661 | 1.33 (0.99) | 1.18 (0.94) | -0.07 ( -0.17, 0.04) | 0.23 |  |
| Heavy rain | 1435 | 1249 | 186 | 1.36 (0.99) | 1.09 (0.86) | **-0.23 (-0.39, -0.07)** | 0.01 | **0.04** |
| No elevated temperature | 5262 | 4661 | 601 | 1.31 (1.00) | 1.16 (0.92) | -0.10 (-0.22, 0.01) | 0.07 |  |
| Elevated temperature | 1830 | 1584 | 246 | 1.41 (0.98) | 1.17 (0.92) | -0.09 (-0.24, 0.05) | 0.20 | 0.58 |
| Below median rain | 3562 | 3189 | 373 | 1.28 (0.97) | 1.17 (0.91) | -0.08 (-0.21, 0.05) | 0.21 |  |
| Above median rain | 3530 | 3056 | 474 | 1.39 (1.01) | 1.15 (0.93) | **-0.14 (-0.26, -0.02)** | 0.02 | **0.10** |
| Below median temperature | 3876 | 3466 | 410 | 1.26 (0.97) | 1.22 (0.97) | -0.03 (-0.15, 0.09) | 0.63 |  |
| Above median temperature | 3216 | 2779 | 437 | 1.43 (1.01) | 1.10 (0.87) | **-0.18 (-0.30, -0.06)** | 0.00 | **0.00** |
| **Caregiver hands** |  |  |  |  |  |  |  |  |
| All observations | 5397 | 4750 | 647 | 1.49 (1.01) | 1.35 (0.93) | 0.01 (-0.10, 0.13) | 0.85 | -- |
| No heavy rain | 4205 | 3705 | 500 | 1.47 (1.00) | 1.32 (0.89) | 0.00 (-0.12, 0.12) | 0.95 |  |
| Heavy rain | 1192 | 1045 | 147 | 1.54 (1.03) | 1.44 (1.02) | 0.05 (-0.15, 0.26) | 0.60 | 0.42 |
| No elevated temperature | 3615 | 3202 | 413 | 1.51 (1.01) | 1.37 (0.89) | 0.03 (-0.10, 0.16) | 0.69 |  |
| Elevated temperature | 1782 | 1548 | 234 | 1.43 (1.00) | 1.31 (0.99) | -0.01 (-0.17, 0.16) | 0.92 | 0.69 |
| Below median rain | 2562 | 2259 | 303 | 1.50 (1.00) | 1.32 (0.88) | -0.04 (-0.19, 0.11) | 0.60 |  |
| Above median rain | 2835 | 2491 | 344 | 1.47 (1.02) | 1.37 (0.96) | 0.06 (-0.08, 0.20) | 0.42 | 0.34 |
| Below median temperature | 2491 | 2201 | 290 | 1.53 (1.00) | 1.41 (0.90) | 0.04 (-0.11, 0.18) | 0.62 |  |
| Above median temperature | 2906 | 2549 | 357 | 1.44 (1.01) | 1.31 (0.94) | -0.01 (-0.15, 0.14) | 0.91 | 0.87 |
| **Soil** |  |  |  |  |  |  |  |  |
| All observations | 2538 | 2264 | 274 | 5.11 (1.06) | 4.96 (1.08) | -0.03 ( -0.20, 0.14) | 0.72 | -- |
| No heavy rain | 1946 | 1742 | 204 | 5.12 (1.10) | 5.00 (1.08) | 0.00 ( -0.21, 0.21) | 1.00 |  |
| Heavy rain | 592 | 522 | 70 | 5.08 (0.92) | 4.86 (1.06) | -0.18 (-0.45, 0.09) | 0.19 | 0.42 |
| No elevated temperature | 2043 | 1824 | 219 | 5.12 (1.05) | 4.96 (1.04) | -0.02 (-0.22, 0.17) | 0.79 |  |
| Elevated temperature | 495 | 440 | 55 | 5.10 (1.11) | 4.96 (1.20) | 0.05 (-0.35, 0.46) | 0.80 | 0.81 |
| Below median rain | 1144 | 1058 | 86 | 5.07 (1.15) | 4.76 (1.25) | -0.16 (-0.49, 0.18) | 0.35 |  |
| Above median rain | 1394 | 1206 | 188 | 5.15 (0.98) | 5.05 (0.97) | 0.00 (-0.18, 0.18) | 1.00 | 0.37 |
| Below median temperature | 1576 | 1438 | 138 | 5.16 (1.06) | 5.03 (1.07) | -0.02 (-0.27, 0.23) | 0.88 |  |
| Above median temperature | 962 | 826 | 136 | 5.03 (1.07) | 4.89 (1.07) | -0.04 (-0.26, 0.19) | 0.75 | 0.92 |
| **Food** |  |  |  |  |  |  |  |  |
| All observations | 2181 | 1945 | 236 | 0.70 (1.40) | 0.63 (1.38) | 0.10 (-0.07, 0.28) | 0.25 | -- |
| No heavy rain | 1685 | 1506 | 179 | 0.49 (1.34) | 0.54 (1.36) | 0.20 (-0.02, 0.41) | 0.07 |  |
| Heavy rain | 496 | 439 | 57 | 1.43 (1.38) | 0.92 (1.42) | -0.32 (-0.70, 0.06) | 0.10 | **0.04** |
| No elevated temperature | 1766 | 1572 | 194 | 0.50 (1.34) | 0.53 (1.37) | 0.16 (-0.05, 0.37) | 0.14 |  |
| Elevated temperature | 415 | 373 | 42 | 1.54 (1.35) | 1.08 (1.37) | -0.20 (-0.63, 0.24) | 0.37 | **0.12** |
| Below median rain | 1020 | 946 | 74 | 0.15 (1.16) | 0.18 (1.25) | 0.18 (-0.09, 0.46) | 0.20 |  |
| Above median rain | 1161 | 999 | 162 | 1.22 (1.42) | 0.84 (1.39) | -0.11 (-0.33, 0.11) | 0.34 | **0.01** |
| Below median temperature | 1382 | 1265 | 117 | 0.33 (1.24) | 0.35 (1.26) | 0.20 (-0.04, 0.44) | 0.11 |  |
| Above median temperature | 799 | 680 | 119 | 1.39 (1.43) | 0.91 (1.44) | -0.13 (-0.42, 0.16) | 0.37 | **0.03** |
| **Flies** |  |  |  |  |  |  |  |  |
| All observations | 610 | 524 | 86 | 2.85 (1.35) | 2.85 (1.34) | -0.05 (-0.40, 0.30) | 0.77 | **--** |
| No heavy rain | 520 | 447 | 73 | 2.83 (1.33) | 2.70 (1.27) | -0.05 (-0.39, 0.30) | 0.78 |  |
| Heavy rain | 90 | 77 | 13 | 3.00 (1.47) | 3.68 (1.40) | -0.04 (-1.14, 1.05) | 0.94 | **0.14** |
| No elevated temperature | 577 | 502 | 75 | 2.84 (1.35) | 2.85 (1.35) | 0.00 (-0.38, 0.38) | 0.99 |  |
| Elevated temperature | 33 | 22 | 11 | 3.26 (1.25) | 2.79 (1.24) | -0.67 (-1.92, 0.59) | 0.30 | 0.25 |
| Below median rain | 320 | 290 | 30 | 2.65 (1.26) | 2.39 (1.19) | -0.05 (-0.58, 0.49) | 0.87 |  |
| Above median rain | 290 | 234 | 56 | 3.11 (1.40) | 3.09 (1.35) | -0.15 (-0.57, 0.28) | 0.50 | 0.44 |
| Below median temperature | 473 | 425 | 48 | 2.77 (1.33) | 2.64 (1.30) | -0.10 (-0.55, 0.36) | 0.67 |  |
| Above median temperature | 137 | 99 | 38 | 3.19 (1.38) | 3.11 (1.34) | **-0.60 (-1.19, -0.01)** | 0.05 | 0.84 |

### **Table S7.** Effect modification by rainfall and temperature within 2 days prior to sampling on adjusted associations between floor material and *E. coli* prevalence in environmental samples. Heavy rain and elevated temperature are each defined as > 80th percentile of daily values during the study period. Above-median rain and temperature are defined as rolling 2-day average values >50^th^ percentile. Interaction p-values <0.20 are denoted in bold and interpreted as evidence of effect modification. The estimates in this table correspond to **Figure S3**.

|  | **N** | | | **Mean prevalence % (n)** | | **Prevalence ratio (95% CI)** | **p-value** | **Interaction**  **p-value** |
| --- | --- | --- | --- | --- | --- | --- | --- | --- |
|  | **Total** | **Unfinished** | **Finished** | **Unfinished** | **Finished** |  |  |  |
| **Stored water** |  |  |  |  |  |  |  |  |
| All observations | 6350 | 5633 | 717 | 76.37 (4302) | 66.53 (477) | 0.95 (0.89, 1.02) | 0.16 | -- |
| No heavy rain | 5085 | 4527 | 558 | 74.30 (3364) | 65.59 (366) | 0.97 (0.90, 1.05) | 0.47 |  |
| Heavy rain | 1265 | 1106 | 159 | 84.81 (938) | 69.81 (111) | **0.89 (0.81, 0.99)** | 0.03 | **0.17** |
| No elevated temperature | 4723 | 4217 | 506 | 74.63 (3147) | 64.03 (324) | 0.94 (0.87, 1.01) | 0.08 |  |
| Elevated temperature | 1627 | 1416 | 211 | 81.57 (1155) | 72.51 (153) | 0.98 (0.89, 1.09) | 0.76 | 0.38 |
| Below median rain | 3222 | 2906 | 316 | 70.54 (2050) | 62.03 (196) | 0.96 (0.87, 1.06) | 0.44 |  |
| Above median rain | 3128 | 2727 | 401 | 82.58 (2252) | 70.07 (281) | **0.92 (0.86, 0.99)** | 0.03 | 0.82 |
| Below median temperature | 3488 | 3140 | 348 | 71.43 (2243) | 65.23 (227) | 0.99 (0.90, 1.08) | 0.79 |  |
| Above median temperature | 2862 | 2493 | 369 | 82.59 (2059) | 67.75 (250) | **0.91 (0.84, 0.98)** | 0.01 | **0.13** |
| **Child hands** |  |  |  |  |  |  |  |  |
| All observations | 7092 | 6245 | 847 | 66.89 (4177) | 58.80 (498) | **0.90 (0.83, 0.98)** | 0.02 | -- |
| No heavy rain | 5657 | 4996 | 661 | 66.49 (3322) | 58.85 (389) | **0.91 (0.83, 1.00)** | 0.05 |  |
| Heavy rain | 1435 | 1249 | 186 | 68.45 (855) | 58.60 (109) | 0.88 (0.76, 1.04) | 0.13 | 0.43 |
| No elevated temperature | 5262 | 4661 | 601 | 65.14 (3036) | 58.57 (352) | **0.91 (0.82, 1.00)** | 0.05 |  |
| Elevated temperature | 1830 | 1584 | 246 | 72.03 (1141) | 59.35 (146) | 0.89 (0.79, 1.01) | 0.07 | 0.41 |
| Below median rain | 3562 | 3189 | 373 | 64.91 (2070) | 58.98 (220) | **0.90 (0.81, 1.00)** | 0.05 |  |
| Above median rain | 3530 | 3056 | 474 | 68.95 (2107) | 58.65 (278) | **0.90 (0.81, 1.00)** | 0.05 | 0.50 |
| Below median temperature | 3876 | 3466 | 410 | 63.65 (2206) | 60.24 (247) | 0.94 (0.84, 1.04) | 0.23 |  |
| Above median temperature | 3216 | 2779 | 437 | 70.92 (1971) | 57.44 (251) | **0.87 (0.78, 0.96)** | 0.01 | **0.03** |
| **Caregiver hands** |  |  |  |  |  |  |  |  |
| All observations | 5397 | 4750 | 647 | 75.96 (3608) | 71.40 (462) | 1.02 (0.95, 1.09) | 0.58 | -- |
| No heavy rain | 4205 | 3705 | 500 | 75.68 (2804) | 71.40 (357) | 1.02 (0.95, 1.09) | 0.63 |  |
| Heavy rain | 1192 | 1045 | 147 | 76.93 (804) | 71.43 (105) | 1.03 (0.89, 1.19) | 0.72 | 0.88 |
| No elevated temperature | 3615 | 3202 | 413 | 77.42 (2479) | 73.61 (304) | 1.03 (0.97, 1.11) | 0.33 |  |
| Elevated temperature | 1782 | 1548 | 234 | 72.93 (1129) | 67.52 (158) | 1.00 (0.88, 1.14) | 0.99 | 0.63 |
| Below median rain | 2562 | 2259 | 303 | 77.69 (1755) | 70.63 (214) | 0.97 (0.88, 1.07) | 0.60 |  |
| Above median rain | 2835 | 2491 | 344 | 74.39 (1853) | 72.09 (248) | 1.06 (0.98, 1.16) | 0.15 | 0.33 |
| Below median temperature | 2491 | 2201 | 290 | 78.42 (1726) | 74.83 (217) | 1.04 (0.96, 1.13) | 0.33 |  |
| Above median temperature | 2906 | 2549 | 357 | 73.83 (1882) | 68.63 (245) | 1.00 (0.91, 1.09) | 1.00 | 0.78 |
| **Soil** |  |  |  |  |  |  |  |  |
| All observations | 2538 | 2264 | 274 | 94.61 (2142) | 93.43 (256) | 0.99 (0.94, 1.03) | 0.50 | -- |
| No heavy rain | 1946 | 1742 | 204 | 93.97 (1637) | 93.13 (190) | 0.99 (0.94, 1.04) | 0.63 |  |
| Heavy rain | 592 | 522 | 70 | 96.74 (505) | 94.29 (66) | 0.97 (0.90, 1.05) | 0.46 | 0.66 |
| No elevated temperature | 2043 | 1824 | 219 | 94.63 (1726) | 93.15 (204) | 0.98 (0.94, 1.03) | 0.51 |  |
| Elevated temperature | 495 | 440 | 55 | 94.54 (416) | 94.55 (52) | 0.98 (0.90, 1.07) | 0.73 | 0.69 |
| Below median rain | 1144 | 1058 | 86 | 92.44 (978) | 86.05 (74) | 0.92 (0.84, 1.03) | 0.15 |  |
| Above median rain | 1394 | 1206 | 188 | 96.52 (1164) | 96.81 (182) | 1.00 (0.97, 1.04) | 0.84 | 0.28 |
| Below median temperature | 1576 | 1438 | 138 | 94.51 (1359) | 92.03 (127) | 0.96 (0.90, 1.02) | 0.16 |  |
| Above median temperature | 962 | 826 | 136 | 94.79 (783) | 94.85 (129) | 0.99 (0.94, 1.04) | 0.73 | 0.54 |
| **Food** |  |  |  |  |  |  |  |  |
| All observations | 2181 | 1945 | 236 | 61.40 (1194) | 58.05 (137) | 1.09 (0.95, 1.25) | 0.20 | -- |
| No heavy rain | 1685 | 1506 | 179 | 57.77 (870) | 58.10 (104) | **1.17 (1.00, 1.38)** | 0.05 |  |
| Heavy rain | 496 | 439 | 57 | 73.80 (324) | 57.89 (33) | 0.84 (0.66, 1.08) | 0.17 | **0.08** |
| No elevated temperature | 1766 | 1572 | 194 | 59.99 (943) | 57.22 (111) | 1.11 (0.94, 1.32) | 0.20 |  |
| Elevated temperature | 415 | 373 | 42 | 67.29 (251) | 61.90 (26) | 1.01 (0.76, 1.34) | 0.96 | 0.98 |
| Below median rain | 1020 | 946 | 74 | 51.59 (488) | 47.30 (35) | 1.13 (0.86, 1.50) | 0.37 |  |
| Above median rain | 1161 | 999 | 162 | 70.67 (706) | 62.96 (102) | 1.00 (0.86, 1.17) | 0.95 | 0.78 |
| Below median temperature | 1382 | 1265 | 117 | 57.94 (733) | 52.99 (62) | 1.07 (0.87, 1.31) | 0.52 |  |
| Above median temperature | 799 | 680 | 119 | 67.79 (461) | 63.03 (75) | 1.07 (0.89, 1.30) | 0.46 | 0.63 |
| **Flies** |  |  |  |  |  |  |  |  |
| All observations | 610 | 524 | 86 | 54.00 (283) | 52.32 (45) | 0.96 (0.75, 1.21) | 0.71 | **--** |
| No heavy rain | 520 | 447 | 73 | 53.91 (241) | 47.94 (35) | 0.92 (0.70, 1.21) | 0.57 |  |
| Heavy rain | 90 | 77 | 13 | 54.54 (42) | 76.92 (10) | 1.09 (0.76, 1.57) | 0.65 | **0.02** |
| No elevated temperature | 577 | 502 | 75 | 52.99 (266) | 52.00 (39) | 0.99 (0.75, 1.29) | 0.91 |  |
| Elevated temperature | 33 | 22 | 11 | 77.27 (17) | 54.55 (6) | **0.53 (0.28, 0.98)** | **0.04** | **0.13** |
| Below median rain | 320 | 290 | 30 | 46.20 (134) | 33.33 (10) | 0.71 (0.41, 1.24) | 0.23 |  |
| Above median rain | 290 | 234 | 56 | 63.68 (149) | 62.50 (35) | 1.05 (0.81, 1.35) | 0.71 | **0.20** |
| Below median temperature | 473 | 425 | 48 | 50.59 (215) | 43.8 (21) | 0.85 (0.58, 1.23) | 0.39 |  |
| Above median temperature | 137 | 99 | 38 | 68.69 (68) | 63.16 (24) | 0.84 (0.65, 1.09) | 0.18 | 0.68 |

### **Table S8.** Effect modification by tertiles of animals owned on adjusted associations between flooring material and log 10-transformed *E. coli* counts in environmental samples. Households owned the following number of animals, mean (range): 1st tertile: 4.6 (0-9), 2nd tertile: 14.4 (10-19), 3rd tertile: 57.6 (20-2700). Interaction p-values <0.20 are denoted in bold and interpreted as evidence of effect modification. The estimates in this table correspond to **Figure 4** in the main text.

|  | **N** | | | **Log10 Mean MPN (SD)** | | **ΔLog10 (95% CI)** | **p-value** | **Interaction**  **p-value** |
| --- | --- | --- | --- | --- | --- | --- | --- | --- |
|  | **Total** | **Unfinished** | **Finished** | **Unfinished** | **Finished** |  |  |  |
| **Stored water** |  |  |  |  |  |  |  |  |
| All observations | 6350 | 5633 | 717 | 0.92 (1.05) | 0.71 (1.05) | -0.01 (-0.10, 0.12) | 0.90 | **--** |
| Animals owned 1^st^ tertile | 2382 | 2087 | 295 | 0.89 (1.05) | 0.75 (1.13) | **0.19 (0.01, 0.38)** | 0.04 |  |
| Animals owned 2^nd^ tertile | 1942 | 1698 | 244 | 0.91 (1.05) | 0.59 (0.95) | -0.04 (-0.21, 0.13) | 0.64 | **0.08** |
| Animals owned 3^rd^ tertile | 2026 | 1848 | 178 | 0.95 (1.05) | 0.79 (1.06) | -0.08 (-0.24, 0.08) | 0.33 | 0.32 |
| **Child hands** |  |  |  |  |  |  |  |  |
| All observations | 7092 | 6245 | 847 | 1.33 (0.99) | 1.16 (0.92) | **-0.10 (-0.20, 0.00)** | 0.05 | **--** |
| Animals owned 1^st^ tertile | 2661 | 2312 | 349 | 1.30 (0.97) | 1.17 (0.93) | 0.09 (-0.08, 0.25) | 0.30 |  |
| Animals owned 2^nd^ tertile | 2156 | 1871 | 285 | 1.35 (1.02) | 1.11 (0.92) | **-0.22 (-0.39, -0.05)** | 0.01 | **0.03** |
| Animals owned 3^rd^ tertile | 2275 | 2062 | 213 | 1.36 (1.00) | 1.21 (0.91) | **-0.16 (-0.32, -0.01)** | 0.04 | 0.29 |
| **Caregiver hands** |  |  |  |  |  |  |  |  |
| All observations | 5397 | 4750 | 647 | 1.49 (1.01) | 1.35 (0.93) | 0.01 (-0.10, 0.13) | 0.85 | **--** |
| Animals owned 1^st^ tertile | 2003 | 1738 | 265 | 1.46 (0.99) | 1.38 (0.95) | **0.22 (0.02, 0.41)** | 0.03 |  |
| Animals owned 2^nd^ tertile | 1650 | 1432 | 218 | 1.49 (1.01) | 1.31 (0.91) | -0.08 (-0.25, 0.10) | 0.39 | **0.20** |
| Animals owned 3^rd^ tertile | 1744 | 1580 | 164 | 1.51 (1.02) | 1.36 (0.91) | -0.09 (-0.29, 0.11) | 0.37 | **0.17** |
| **Soil** |  |  |  |  |  |  |  |  |
| All observations | 2538 | 2264 | 274 | 5.11 (1.06) | 4.96 (1.08) | -0.03 ( -0.20, 0.14) | 0.72 | **--** |
| Animals owned 1^st^ tertile | 957 | 843 | 114 | 5.08 (1.04)) | 4.87 (1.13) | 0.02 (-0.24, 0.27) | 0.91 |  |
| Animals owned 2^nd^ tertile | 767 | 674 | 93 | 5.10 (1.08) | 4.88 (1.03) | -0.02 (-0.23, 0.18) | 0.80 | 0.79 |
| Animals owned 3^rd^ tertile | 814 | 747 | 67 | 5.17 (1.07) | 5.23 (1.02) | 0.06 (-0.17, 0.28) | 0.63 | **0.12** |
| **Food** |  |  |  |  |  |  |  |  |
| All observations | 2181 | 1945 | 236 | 0.70 (1.40) | 0.63 (1.38) | 0.10 (-0.07, 0.28) | 0.25 | **--** |
| Animals owned 1^st^ tertile | 831 | 727 | 104 | 0.67 (1.38) | 0.80 (1.46) | 0.23 (-0.08, 0.54) | 0.15 |  |
| Animals owned 2^nd^ tertile | 668 | 588 | 80 | 0.73 (1.42) | 0.44 (1.30) | -0.11 (-0.46, 0.24) | 0.54 | **0.04** |
| Animals owned 3^rd^ tertile | 682 | 630 | 52 | 0.71 (1.41) | 0.58 (1.35) | 0.18 (-0.23, 0.58) | 0.39 | 0.61 |
| **Flies** |  |  |  |  |  |  |  |  |
| All observations | 610 | 524 | 86 | 2.85 (1.35) | 2.85 (1.34) | -0.05 (-0.40, 0.30) | 0.77 | **--** |
| Animals owned 1^st^ tertile | 243 | 202 | 41 | 2.83 (1.33) | 2.76 (1.30) | -0.02 (-0.53, 0.49) | 0.93 |  |
| Animals owned 2^nd^ tertile | 189 | 167 | 22 | 2.78 (1.32) | 2.99 (1.37) | 0.25 (-0.42, 0.92) | 0.46 | 0.34 |
| Animals owned 3^rd^ tertile | 178 | 155 | 23 | 2.97 (1.40) | 2.87 (1.45) | -0.12 (-0.82, 0.59) | 0.75 | 0.90 |

### **Table S9.** Effect modification by tertiles of animals owned on adjusted associations between floor material and *E. coli* prevalence in environmental samples. Households owned the following number of animals, mean (range): 1st tertile: 4.6 (0-9), 2^nd^ tertile: 14.4 (10-19), 3^rd^ tertile: 57.6 (20-2700). Interaction p-values <0.20 are denoted in bold and interpreted as evidence of effect modification. The estimates in this table correspond to **Figure S4**.

|  | **N** | | | **Mean prevalence % (n)** | | **Prevalence ratio (95% CI)** | **p-value** | **Interaction**  **p-value** |
| --- | --- | --- | --- | --- | --- | --- | --- | --- |
|  | **Total** | **Unfinished** | **Finished** | **Unfinished** | **Finished** |  |  |  |
| **Stored water** |  |  |  |  |  |  |  |  |
| All observations | 6350 | 5633 | 717 | 76.37 (4302) | 66.53 (477) | 0.95 (0.89, 1.02) | 0.16 | -- |
| Animals owned 1^st^ tertile | 2382 | 2087 | 295 | 75.37 (1573) | 65.76 (194) | 1.01 (0.89, 1.14) | 0.84 |  |
| Animals owned 2^nd^ tertile | 1942 | 1698 | 244 | 75.8 (1287) | 65.57 (160) | 0.98 (0.87, 1.11) | 0.75 | 0.97 |
| Animals owned 3^rd^ tertile | 2026 | 1848 | 178 | 78.03 (1442) | 69.1 (123) | 0.91 (0.82, 1.00) | 0.04 | 0.76 |
| **Child hands** |  |  |  |  |  |  |  |  |
| All observations | 7092 | 6245 | 847 | 66.89 (4177) | 58.80 (498) | **0.90 (0.83, 0.98)** | 0.02 | -- |
| Animals owned 1^st^ tertile | 2661 | 2312 | 349 | 66.18 (1530) | 59.89 (209) | 1.02 (0.90, 1.15) | 0.73 |  |
| Animals owned 2^nd^ tertile | 2156 | 1871 | 285 | 66.76 (1249) | 53.68 (153) | **0.81 (0.69, 0.95)** | 0.01 | **0.09** |
| Animals owned 3^rd^ tertile | 2275 | 2062 | 213 | 67.8 (1398) | 63.85 (136) | 0.91 (0.80, 1.05) | 0.21 | 0.83 |
| **Caregiver hands** |  |  |  |  |  |  |  |  |
| All observations | 5397 | 4750 | 647 | 75.96 (3608) | 71.40 (462) | 1.02 (0.95, 1.09) | 0.58 | -- |
| Animals owned 1^st^ tertile | 2003 | 1738 | 265 | 75.14 (1306) | 71.32 (189) | 1.10 (0.96, 1.25) | 0.17 |  |
| Animals owned 2^nd^ tertile | 1650 | 1432 | 218 | 75.42 (1080) | 70.18 (153) | 1.00 (0.90, 1.11) | 0.97 | 0.45 |
| Animals owned 3^rd^ tertile | 1744 | 1580 | 164 | 77.34 (1222) | 73.17 (120) | 0.99 (0.87, 1.13) | 0.90 | 0.54 |
| **Soil** |  |  |  |  |  |  |  |  |
| All observations | 2538 | 2264 | 274 | 94.61 (2142) | 93.43 (256) | 0.99 (0.94, 1.03) | 0.50 | -- |
| Animals owned 1^st^ tertile | 957 | 843 | 114 | 95.02 (801) | 91.23 (104) | 0.97 (0.91, 1.04) | 0.36 |  |
| Animals owned 2^nd^ tertile | 767 | 674 | 93 | 94.21 (635) | 94.62 (88) | 1.02 (0.96, 1.08) | 0.47 | **0.14** |
| Animals owned 3^rd^ tertile | 814 | 747 | 67 | 94.51 (706) | 95.52 (64) | 0.99 (0.93, 1.05) | 0.71 | 0.24 |
| **Food** |  |  |  |  |  |  |  |  |
| All observations | 2181 | 1945 | 236 | 61.40 (1194) | 58.05 (137) | 1.09 (0.95, 1.25) | 0.20 | -- |
| Animals owned 1^st^ tertile | 831 | 727 | 104 | 60.66 (441) | 63.46 (66) | 1.15 (0.92, 1.43) | 0.21 |  |
| Animals owned 2^nd^ tertile | 668 | 588 | 80 | 62.41 (367) | 51.25 (41) | 1.01 (0.77, 1.32) | 0.95 | **0.09** |
| Animals owned 3^rd^ tertile | 682 | 630 | 52 | 61.27 (386) | 57.69 (30) | 1.10 (0.84, 1.44) | 0.49 | 0.59 |
| **Flies** |  |  |  |  |  |  |  |  |
| All observations | 610 | 524 | 86 | 54.00 (283) | 52.32 (45) | 0.96 (0.75, 1.21) | 0.71 | -- |
| Animals owned 1^st^ tertile | 243 | 202 | 41 | 52.48 (106) | 53.66 (22) | 1.17 (0.81, 1.67) | 0.40 |  |
| Animals owned 2^nd^ tertile | 189 | 167 | 22 | 54.49 (91) | 54.55 (12) | 1.04 (0.67, 1.56) | 0.87 | 0.86 |
| Animals owned 3^rd^ tertile | 178 | 155 | 23 | 55.48 (86) | 47.83 (11) | 0.84 (0.51, 1.38) | 0.50 | 0.42 |
